# SUMOylation targets shugoshin to stabilize sister kinetochore biorientation

**DOI:** 10.1101/840157

**Authors:** Xue Bessie Su, Menglu Wang, Claudia Schaffner, Dean Clift, Olga O. Nerusheva, Triin Tammasalu, Andreas Wallek, Yehui Wu, David A. Kelly, A. Arockia Jeyaprakash, Zuzana Storchova, Ronald Hay, Adèle L. Marston

## Abstract

The accurate segregation of chromosomes during mitosis relies on the attachment of sister chromatids to microtubules from opposite poles, called biorientation. Sister chromatid cohesion resists microtubule forces, generating tension which provides the signal that biorientation has occurred. How tension silences the surveillance pathways that prevent cell cycle progression and correct erroneous kinetochore-microtubule remains unclear. Here we identify SUMOylation as a mechanism that promotes anaphase onset upon biorientation. SUMO ligases modify the tension-sensing pericentromere-localized chromatin protein, shugoshin, to stabilize bioriented sister kinetochore-microtubule attachments. In the absence of SUMOylation, Aurora B kinase removal from kinetochores is delayed. Shugoshin SUMOylation prevents its binding to protein phosphatase 2A (PP2A) and release of this interaction is important for stabilizing sister kinetochore biorientation. We propose that SUMOylation modulates the kinase-phosphatase balance within pericentromeres to inactivate the error correction machinery, thereby allowing anaphase entry in response to biorientation.

## Introduction

Mitosis divides the nucleus to produce two genetically identical daughter cells. Prior to mitosis, DNA replication produces sister chromatids, linked together by the cohesin complex. Sister chromatids are aligned at metaphase, thus allowing microtubule spindles to be captured by kinetochores assembled on centromeres. The correct form of attachment is termed ‘biorientation’, meaning that the kinetochores on the two sister chromatids are attached to microtubules emanating from opposite spindle poles. Biorientation creates tension, because cohesin holding sister chromatids together resists the pulling force of microtubules [1]. The fulfilment of biorientation allows securin degradation and, consequently, the activation of the protease separase, which cleaves cohesin, triggering sister chromatid separation (reviewed in [2]).

The conserved shugoshin protein plays key roles in promoting biorientation in mitosis and preventing cell cycle progression where biorientation fails [3, 4]. Budding yeast possesses a single shugoshin gene, *SGO1*. Sgo1 localizes to both the core ∼125bp centromere, where the kinetochore resides, and the surrounding ∼20kb cohesin-rich chromosomal region called the pericentromere [5]. The kinetochore-localized Bub1 kinase promotes Sgo1 enrichment at the pericentromere through phosphorylation of S121 on histone H2A [6–8]. Sgo1, in turn, recruits condensin and protein phosphatase 2A, PP2A-Rts1, to the pericentromere and maintains the chromosome passenger complex (CPC) containing Aurora B kinase at centromeres during mitosis [9, 10]. Condensin at pericentromeres is thought to bias the conformation of the sister chromatids to favour biorientation. The CPC recognizes erroneous microtubule-kinetochore attachments and destabilizes them, thereby maintaining the activity of the spindle assembly checkpoint (SAC) to prevent anaphase entry (reviewed in [11]). In vertebrate cells, PP2A-B56 protects cohesin in pericentromeres from removal by the so-called prophase pathway, which removes cohesin through a non-proteolytic mechanism that is independent of separase [12, 13]. In budding yeast, PP2A-Rts1 is recruited by shugoshin despite the absence of the prophase pathway [9,10,14]. Instead, PP2A-Rts1 has been implicated in ensuring the equal segregation of sister chromatids during mitosis, since mutants failing to recruit PP2A-Rts1 to the centromere are unable to respond to a lack of inter-sister kinetochore tension and mis-segregate chromosomes upon recovery [9, 14].

Sgo1 both directs and responds to cell cycle cues as chromosomes establish and achieve biorientation, upon which anaphase entry is triggered. A key signal that biorientation has occurred is the tension-dependent removal of Sgo1 from the pericentromere during metaphase, resulting in the delocalization of its effectors, PP2A-Rts1, condensin and the CPC [8, 14]. Upon anaphase I onset, Sgo1 is ubiquitinated and degraded by APC/C-Cdc20 [14, 15]. Similarly, human shugoshin is degraded in anaphase as a result of APC/C-Cdc20 activity [16]. Shugoshin can be stabilized by mutation of its APC-Cdc20-dependent destruction sequence in both yeast and human cells, however, this does not impair the metaphase-anaphase transition [13,14,17]. Nevertheless, there is evidence that Sgo1 inactivation is important, since *SGO1* overexpression results in a pronounced metaphase delay and a block to cohesin cleavage [18]. Since the Sgo1-induced metaphase delay is abrogated by deletion of *BUB1*, it is likely that Sgo1 must be localized at the pericentromere to prevent anaphase onset [18]. Accordingly, we showed that Sgo1 and its associated proteins are released from centromeres upon sister chromatid biorientation and tension [8]. Potentially, the temporal separation between Sgo1 removal from each pericentromere and its later degradation could ensure the reversibility of the mechanism that monitors biorientation right up until the moment that the commitment to anaphase is made. However, this model poses a conundrum: with the initiation of cohesin cleavage at anaphase onset, tension between sister kinetochores is lost, which could lead to re-activation of the error correction and biorientation pathways. What is more, it was unclear whether Sgo1-associated PP2A-Rts1 and CPC depart from centromeres simply by interacting with Sgo1, or by a more sophisticated mechanism involving finely orchestrated interplay between kinase and phosphatase activities.

Here, we identify Sgo1 SUMOylation as a mechanism that inactivates the pericentromeric signalling hub and thereby ensures timely anaphase onset. Small ubiquitin-like modifier (SUMO) is a 12 kDa protein that is covalently added to lysine residues of SUMO substrates. SUMOylation is carried out by the sequential activities of E1 activation enzyme, E2 conjugation enzyme and E3 ligase [19]. We isolated the E3 SUMO ligase, Siz2, as a negative regulator of Sgo1 through a genetic screen. We show that Sgo1 is SUMOylated in a manner dependent on Siz2 together with its paralog, Siz1, and that a failure to SUMOylate Sgo1 causes a delay to anaphase onset. We provide evidence that Sgo1 SUMOylation modulates its interaction with PP2A-Rts1 and facilitates the removal of CPC to stabilize sister chromatid biorientation, thereby enabling efficient entry into anaphase.

## Results

### SUMO ligases reverse the effects of SGO1 overexpression

To identify negative regulators of Sgo1, we screened for high copy suppressors of the poor growth caused by *SGO1* overexpression [18]. We recovered a number of plasmids that improved the growth of cells carrying multiple copies of *SGO1* under galactose-inducible control (*pGAL-SGO1*) (Figure S1A; Table S1), including one carrying a ∼5kb fragment containing truncated *SLP1* and *PUP1* together with full length *ISN1* and *SIZ2* (Figure 1A). *SIZ2*, encoding one of three budding yeast SUMO E3 ligases and sharing functional redundancy with its paralog, Siz1, is an attractive candidate for an Sgo1 antagonist, since Siz1/Siz2 have functions in chromosome segregation and cell division (Johnson and Gupta, 2001; Makhnevych et al., 2009; Montpetit et al., 2006; Takahashi et al., 2006). We used live cell imaging to confirm that overexpression of *SIZ2* (by placement under control of the copper-inducible promoter, *pCUP1-SIZ2)* counteracts the delay in metaphase caused by *SGO1* overexpression (Figure 1B). Metaphase duration, estimated as the time between spindle pole body separation (emergence of two Spc42-tdTomato foci) and anaphase onset (dispersal of Cdc14-GFP from the nucleolus), was markedly reduced in *pGAL-SGO1 pCUP1-SIZ2* cells compared to *pGAL-SGO1* cells (Figure 1C). These findings suggest that Siz1/Siz2 promote anaphase onset by counteracting Sgo1. Indeed, *siz1*Δ *siz2*Δ cells show a prolonged metaphase, measured as the time between formation of a short metaphase spindle (Tub1-YFP) and Cdc14-GFP dispersal (Figure 1D and E; [20]). Consistently, short metaphase spindles and securin (Pds1) persist in *siz1*Δ *siz2*Δ cells after synchronous release from a G1 arrest (Figure 1F and 1G). Deletion of *SGO1* in *siz1*Δ *siz2*Δ cells partially rescued the metaphase delay, as judged by both decreased accumulation of short spindles and reduced Pds1 stabilization (Figure 1F and 1G), as did deletion of *CDC55* (Figure S1B), previously shown to rescue the metaphase delay of *pGAL-SGO1*[18], further indicating that ectopic Sgo1 activity is at least partially responsible for the metaphase delay of cells lacking *SIZ1* and *SIZ2*. We note, however, that the metaphase delay was more pronounced in *sgo1*Δ *siz1*Δ *siz2*Δ cells than in *sgo1*Δ cells, indicating that additional factors prevent timely anaphase onset in the absence of Siz1-Siz2. We conclude that Siz1/Siz2 promote anaphase onset, in part by antagonising Sgo1.

**Figure 1.**
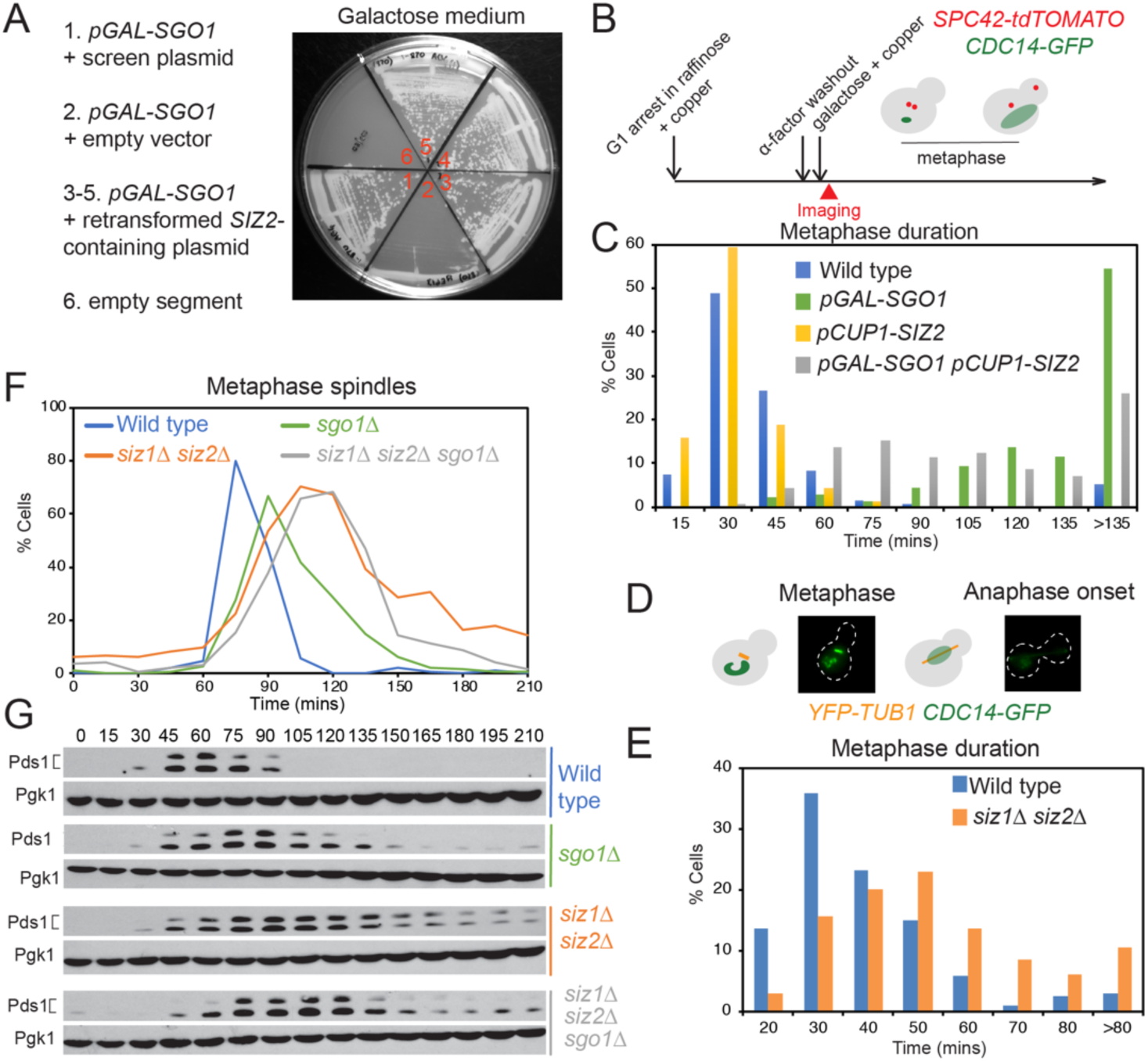
SUMO ligases promote Sgo1 inactivation to allow timely anaphase progression. (A) Overexpression of *SIZ2* rescues the slow growth phenotype of *SGO1* overexpression. A *pGAL-SGO1* (AMy870) strain carrying empty vector (AMp67), or with yEP13-(*SLP1*) *ISN1 SIZ2* (*PUP1*) (AMp1435) was streaked onto medium containing galactose. (B and C) *SIZ2* overexpression partially rescues the metaphase delay of *SGO1-*overexpressing cells. (B) Schematic of live cell imaging experiment. Cells carrying Spc42-tdTomato and Cdc14-GFP were synchronized in G1 in media containing 2% raffinose. 25 μM copper sulfate was added to induce *pCUP1-SIZ2* expression. After releasing from G1, 0.2% galactose was added to induce *pGAL-SGO1* expression. The duration of metaphase was estimated by the time taken between the separation of the spindle pole bodies (two Spc42-tdTomato foci) and the dispersal of Cdc14-GFP from the nucleolus. (C) Metaphase duration is shown for wild type (AMy24115), *pGAL-SGO1* (AMy27596), *pGAL-SGO1 pCUP1-SIZ2* (AMy27738) and *pCUP1-SIZ2* (AMy27952) strains. (D and E) Siz1 and Siz2 are required for timely anaphase onset. Metaphase duration was determined as the time between formation of a short bipolar spindle (YFP-Tub1) and release of Cdc14-GFP from the nucleolus from live cell imaging. (D) Schematics and representative images are shown. (E) Metaphase duration is shown for wild type (AMy24174) and *siz1*Δ *siz2*Δ (AMy24313) strains. (F) and (G) The metaphase delay of *siz1*Δ *siz2*Δ cells is partially rescued by *SGO1* deletion. Wild type (AMy1290), *sgo1*Δ (AMy8466), *siz1*Δ *siz2*Δ (AMy8465) and *siz1*Δ *siz2*Δ *sgo1*Δ (AMy12110) strains carrying *PDS1-6HA* were released from a G1 arrest. Spindle morphology was scored after anti-tubulin immunofluorescence and the percentages of short (metaphase) spindles are shown (top) and Pds1 levels were analysed by anti-HA Western blot (bottom). Pgk1 is shown as a loading control.

### A chromatin-associated pool of Sgo1 is SUMOylated by Siz1 and/or Siz2

To determine whether Siz1/Siz2 might counteract Sgo1 by direct SUMOylation, we assayed Sgo1-SUMO conjugates *in vivo*. Cells carrying *SGO1-6HA* and a plasmid producing His-tagged yeast SUMO (7His-Smt3) were lysed under denaturing conditions and SUMOylated proteins were isolated using nickel affinity chromatography (Figure 2A). Anti-HA western blotting identified two slow-migrating bands corresponding to SUMOylated Sgo1 in wild type but not *siz1*Δ *siz2*Δ cells (Figure 2B). We further demonstrated direct SUMOylation of purified Sgo1 (Figure S2) *in vitro* by either Siz1 or Siz2 (Figure 2C).

**Figure 2.**
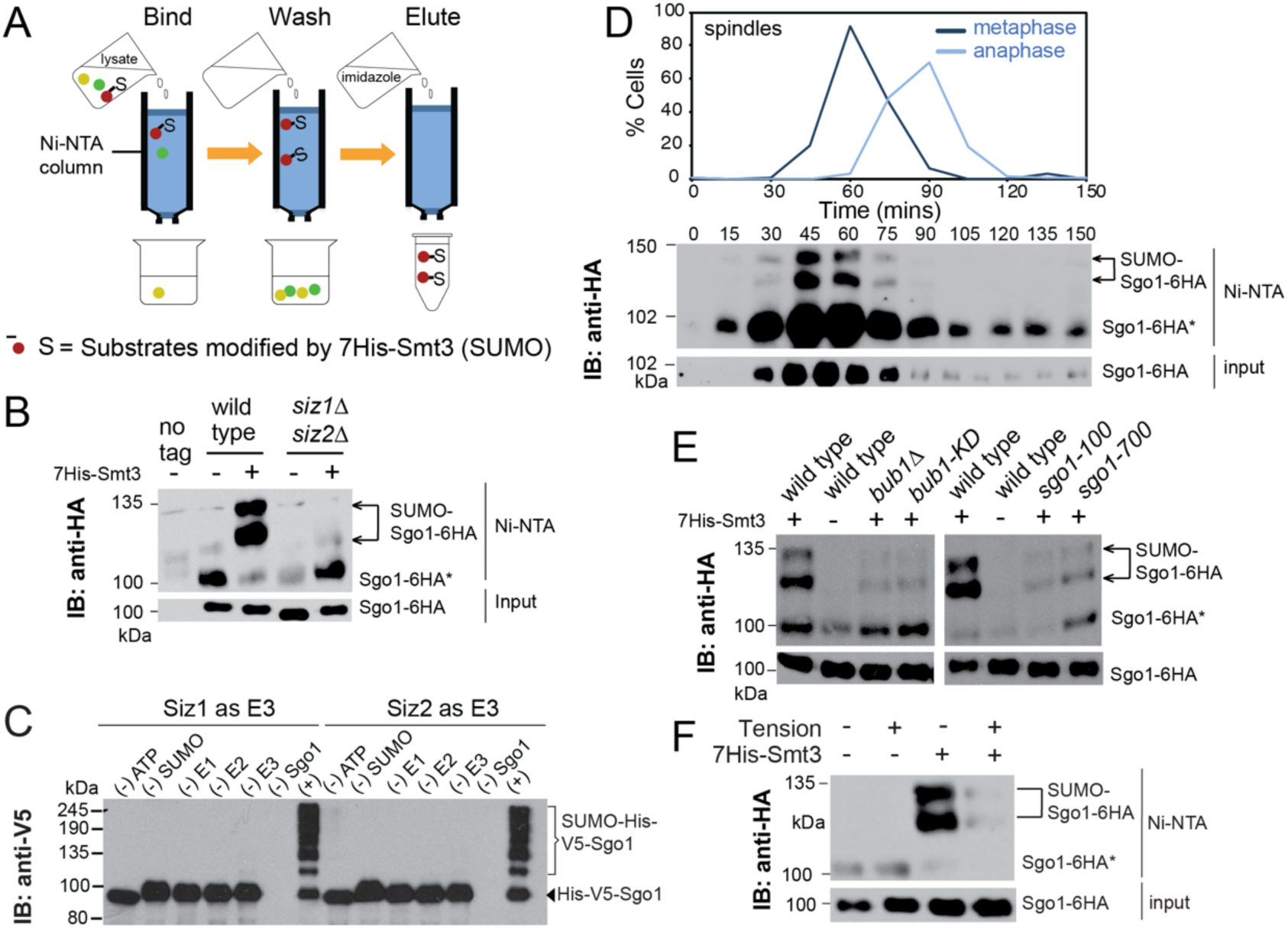
Sgo1 is SUMOylated, depending on its association with pericentromeres. (A and B) Sgo1 is SUMOylated in a Siz1/Siz2-dependent manner. (A) Scheme describing purification of SUMOylated proteins. (B) Extracts from untagged (AMy1176), *SGO1-6HA* (AMy906) and *siz1*Δ *siz2*Δ *SGO1-6HA* (AMy7911) strains carrying empty vector (pRS426), or with *7*×*HIS-SMT3* (AMp773) were purified over Ni-NTA resin and anti-HA immunoblot was performed on both input and elute. Arrows and asterisks indicate SUMO-Sgo1-6HA and unmodified Sgo1-6HA, which binds non-specifically to the resin, respectively. (C) Sgo1 is SUMOylated by Siz1 and Siz2 *in vitro*. Purified Sgo1 was incubated with 1 μM E1, E2, E3, SUMO and ATP or missing one component as indicated. Reaction was incubated at 30°C for 3 h. (D) Sgo1 SUMOylation occurs in metaphase. Cells carrying *SGO1-6HA* and *7xHIS-SMT3* (AMy7655) were released from G1, harvested at the indicated intervals, and SUMOylation was analysed as described in (A). Cell cycle stage was monitored by scoring spindle morphology after anti-tubulin immunofluorescence. (E) Chromatin association promotes Sgo1 SUMOylation. Sgo1 SUMOylation was determined in wild type (AMy7654), *bub1*Δ (AMy10098), *bub1-KD* (catalytically inactive Bub1 kinase, AMy10102), *sgo1-100* (AMy26334) and *sgo1-700* (AMy26336) strains. (F) Sgo1 SUMOylation is lost upon the establishment of tension between sister kinetochores. Cells carrying *pMET-CDC20* and either *7xHIS-SMT3* (AM9641) or empty vector (AMy26342) were arrested in metaphase by depletion of Cdc20 either in the presence of benomyl and nocodazole (no tension) or DMSO (tension).

Sgo1 is recruited to the pericentromeric chromatin during S phase, released into the nucleoplasm upon sister kinetochore biorientation at metaphase and degraded in anaphase [8]. Sgo1 SUMOylation was maximal prior to anaphase onset, reflecting Sgo1 abundance (Figure 2D). We assessed whether Sgo1 recruitment to pericentromeric chromatin is important for its SUMOylation. Loss of Bub1 or inactivation of its kinase activity which is required for phosphorylation of histone H2A-S121 and Sgo1 recruitment to pericentromeres [6, 21] greatly diminished Sgo1 SUMOylation (Figure 2E). Similarly, the *sgo1-100* and *sgo1-700* alleles, which carry point mutations that delocalize Sgo1 from pericentromeres [10], also markedly reduced Sgo1 SUMOylation (Figure 2E). To test whether Sgo1 removal from pericentromeres in response to tension coincides with loss of SUMOylation, we analysed cells arrested in metaphase (by depletion of *CDC20*) either in the presence (-tension) or absence (+tension) of microtubule depolymerising drugs. Indeed, spindle tension largely abrogated Sgo1 SUMOylation, though total Sgo1 levels were comparable to the no tension condition (Figure 2F). We conclude that Sgo1 SUMOylation is promoted by its association with pericentromeres.

### Sgo1 SUMOylation requires its coiled-coil

If Sgo1 SUMOylation mediates the effect of Siz1/Siz2 in the metaphase-anaphase transition, a mutant that specifically abrogates Sgo1 SUMOylation, either by losing the SUMOylation sites, or by reducing Sgo1’s interaction with the E3 ligases, is expected to recapitulate *siz1*Δ *siz2*Δ’s metaphase delay. With the aim to map the SUMOylation sites, we performed mass spectrometry analysis of elutes from nickel affinity chromatography identified Lys124 as an *in vivo* SUMOylation site (Table S2). However, the *sgo1-K124R* mutant retained high levels of SUMOylation, suggesting the presence of additional SUMOylation sites (Figure 3A and B). As an alternative approach to mass spectrometry, we analysed a series of Sgo1 truncations (Figure S3A). SUMOylation was abolished in Sgo1-Δ2-208 and reduced in Sgo1-Δ2-108, suggesting that the N-terminal 108 amino acids are important for SUMOylation together with K124R (Figure S3B). Further analysis revealed robust SUMOylation of Sgo1-Δ2-40 and Sgo1-Δ2-40-K124R but not Sgo1-Δ2-108 K124R (Figure S3B), suggesting that SUMOylation requires amino acids 40-108. Interestingly, this region encompasses the coiled-coil domain of budding yeast Sgo1, which is known to directly bind PP2A-Rts1 and is also important for maintaining CPC at the centromeres (Verzijlbergen et al., 2014; Xu et al., 2009). We mutated the Lys residues in this region in several combinations (Figure 3A). SUMOylation was progressively reduced in the Sgo1-K56R K85R (Sgo1-2R), Sgo1-K56R K64R K70R K85R (Sgo1-4R) and Sgo1-K56R K64R K70R K85R K124R (Sgo1-5R) mutants (Figure 3B). The reduced SUMOylation of Sgo1-4R was recapitulated in the *in vitro* SUMOylation assay (Figure 3C). Therefore, lysines in Sgo1 coiled-coil enable its SUMOylation and may be direct targets of Siz1/Siz2. Moreover, mutation of these lysines specifically reduced SUMOylation in Sgo1 both *in vivo* and *in vitro*, providing a tool to study the effects of reduced Sgo1 SUMOylation *in vivo*.

**Figure 3.**
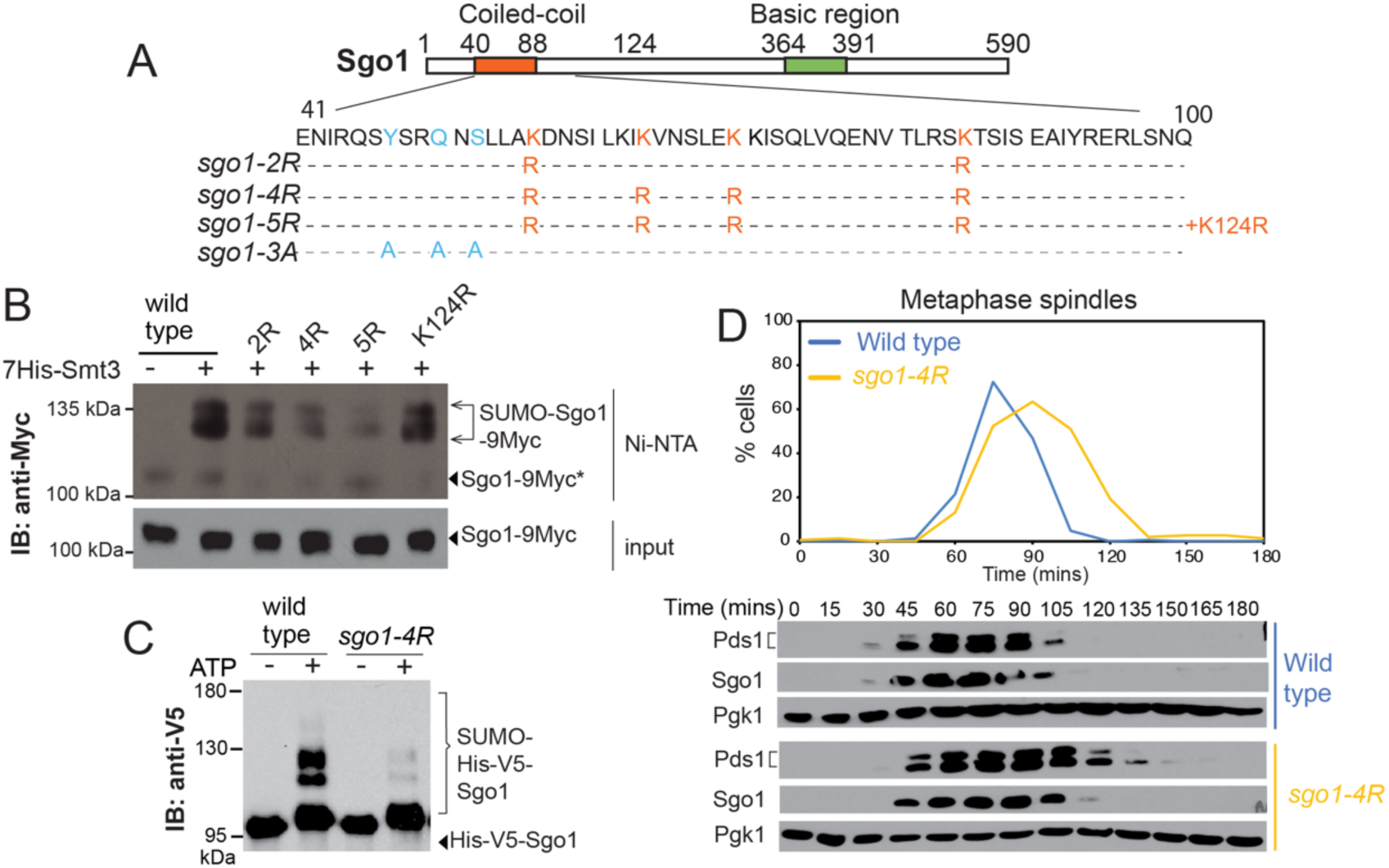
Sgo SUMOylation requires residues within its coiled-coil and is important for timely anaphase onset. (A) Schematic of Sgo1 showing the sequence of the coiled-coil domain (bottom) and residues mutated in the indicated mutants. (B) Sgo1 SUMOylation requires residues K56, K64, K70, K85 and K124. Strains for *in vivo* SUMOylation analysis carried Sgo1-9Myc and were wild type (AMy24367), *sgo1-K56R K85R* (‘2R’, AMy24299), *sgo1-K56R K64R K70R K85R* (‘4R’, AMy23828), *sgo1-K56R K64R K70R K85R K124R* (‘5R’, AMy24371) and *sgo1-K124R* (AMy24369). (C) SUMOylation is reduced *in vitro* for the *sgo1-4R* mutant. Purified Sgo1 and Sgo1-4R proteins were SUMOylated *in vitro* using 0.1 μM E1-E3, in the presence or absence of ATP. (D) The *sgo1-4R* mutant is delayed in metaphase. Cell cycle analysis of wild type (AMy8467) and *sgo1-K56R K64R K70R K85R*-9Myc (AMy23934) strains carrying *SGO1-9MYC* and *PDS1-3HA* was performed as described in Figure 1F.

Time course analysis and live cell imaging revealed a negligible or mild metaphase delay in the *sgo1-K124R* and *sgo1-2R* mutants, respectively (Figure S3C and D) and the *sgo1-2R* mutation did not exacerbate the metaphase delay of *siz1*Δ *siz2*Δ cells (Figure S3E). The *sgo1-4R* mutant showed a more pronounced metaphase delay (Figure 3D), but additional mutation of K124R (*sgo1-5R*) largely abrogated this delay (Figure S3F), suggesting adverse effects on Sgo1 protein function (see also Figure 5B below). Therefore, we focused on *sgo1-4R* for further analysis of the effects of reduced Sgo1 SUMOylation.

**Figure 5.**
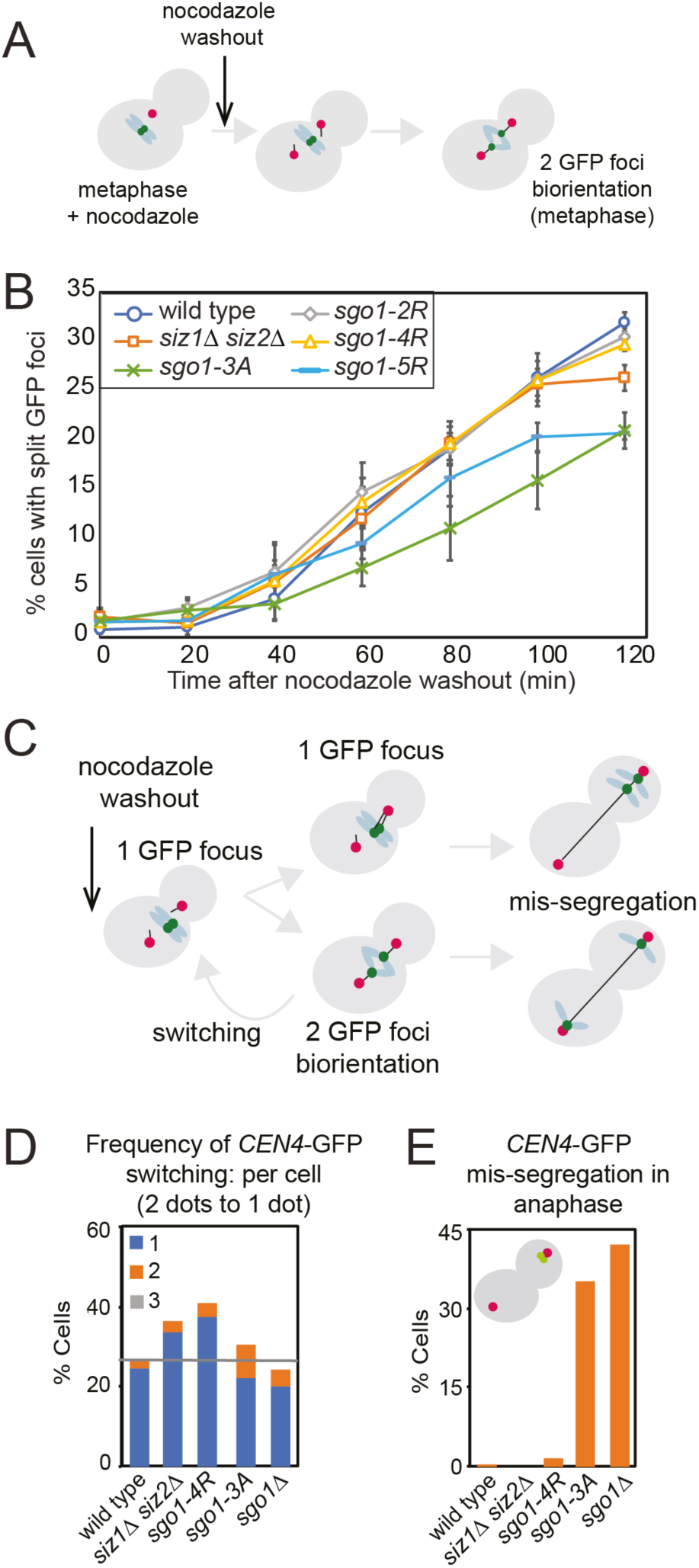
SUMOylation of Sgo1 facilitates stable microtubule-kinetochore attachments. (A and B) UnSUMOylated Sgo1 is proficient for sister kinetochore biorientation. (A) Experimental scheme for evaluating sister kinetochore biorientation after nocodazole washout. Briefly, cells were arrested in metaphase in the presence of nocodazole. After drug washout, cells remained arrested in metaphase and were fixed and visualized as described in ‘Methods’. (B) In contrast to the *sgo1-3A* mutant, *sgo1-2R*,-*4R* and *siz1*Δ *siz2*Δ cells are proficient in the initial establishment of sister kinetochore biorientation, though *sgo1-5R* is partially defective in biorientation. Strains used carried *pMET-CDC20*, *CEN4-GFP* and *SPC42-dtTOMATO*, and were wild type (AMy4643), *sgo1-3A* (AMy8923), *siz1*Δ *siz2*Δ (AMy26844), *sgo1-2R* (AMy26753), *sgo1-4R* (AMy26278) and *sgo1-5R* (AMy26758). The number of cells with two *CEN4-GFP* foci was determined for 200 cells. (C-E) Unstable kinetochore-microtubule attachments in SUMOylation mutants. (C) Scheme describing biorientation assay after nocodazole washout. All strains were imaged for a total of 3 h post nocodazole washout. (D) Biorientation is unstable in *siz1*Δ *siz2*Δ and *sgo1-4R* cells. Shown are the percentages of cells with the indicated number of switching(s) from two *CEN4-GFP* foci back to one focus. (E) Unlike *sgo1*Δ and *sgo1-3A*, *siz1*Δ *siz2*Δ and *sgo1-4R* cells do not show increased mis-segregation of chromosomes. The inheritance of *CEN4-GFP* by the daughter cells was analyzed for each strain. Strains used in (C-E) contained *CEN4-GFP*, *pMET-CDC20* and *SPC42-tdTOMATO* and were wild type (AMy4643), *sgo1-3A* (AMy8923), *sgo1*Δ (AMy6117), *siz1*Δ *siz2*Δ (AMy26844) and *sgo1-4R* (AMy26278). Typically >100 cells and at least 50 cells were analyzed for each strain. at each time point. Error bars represent standard error of 3 biological replicates.

### Sgo1 is not a direct target of Slx5-Slx8-mediated ubiquitination

How might SUMOylation of Sgo1 reverse its inhibitory effects on anaphase onset? Sgo1 was stabilized in cells lacking E3 ligases, Siz1 and Siz2 (Figure S1B), or where E2 Ubc9 function was impaired (Figure S4A). Cells lacking the kinetochore-associated Slx5/Slx8 SUMO-targeted ubiquitin ligase (STUbL) complex show increased centromeric PP2A-Rts1 [22] and *slx5*Δ cells exhibit a metaphase delay in which Sgo1 is stabilized (Figure 4A). This led us to test whether Sgo1-SUMO conjugates are targeted for proteosomal degradation by Slx5/8 [23]. We measured the half-life of Sgo1 protein levels after transient expression of ectopic *pGAL-SGO1* in G1-arrested cells followed by quenching with glucose/cycloheximide (Figure S4B). This revealed a partial stabilization of Sgo1 in both *siz1*Δ *siz2*Δ and *slx5*Δ, compared to wild type cells (Figure S4C), indicating that Siz1/Siz2 and Slx5/Slx8 promote Sgo1 degradation.

**Figure 4.**
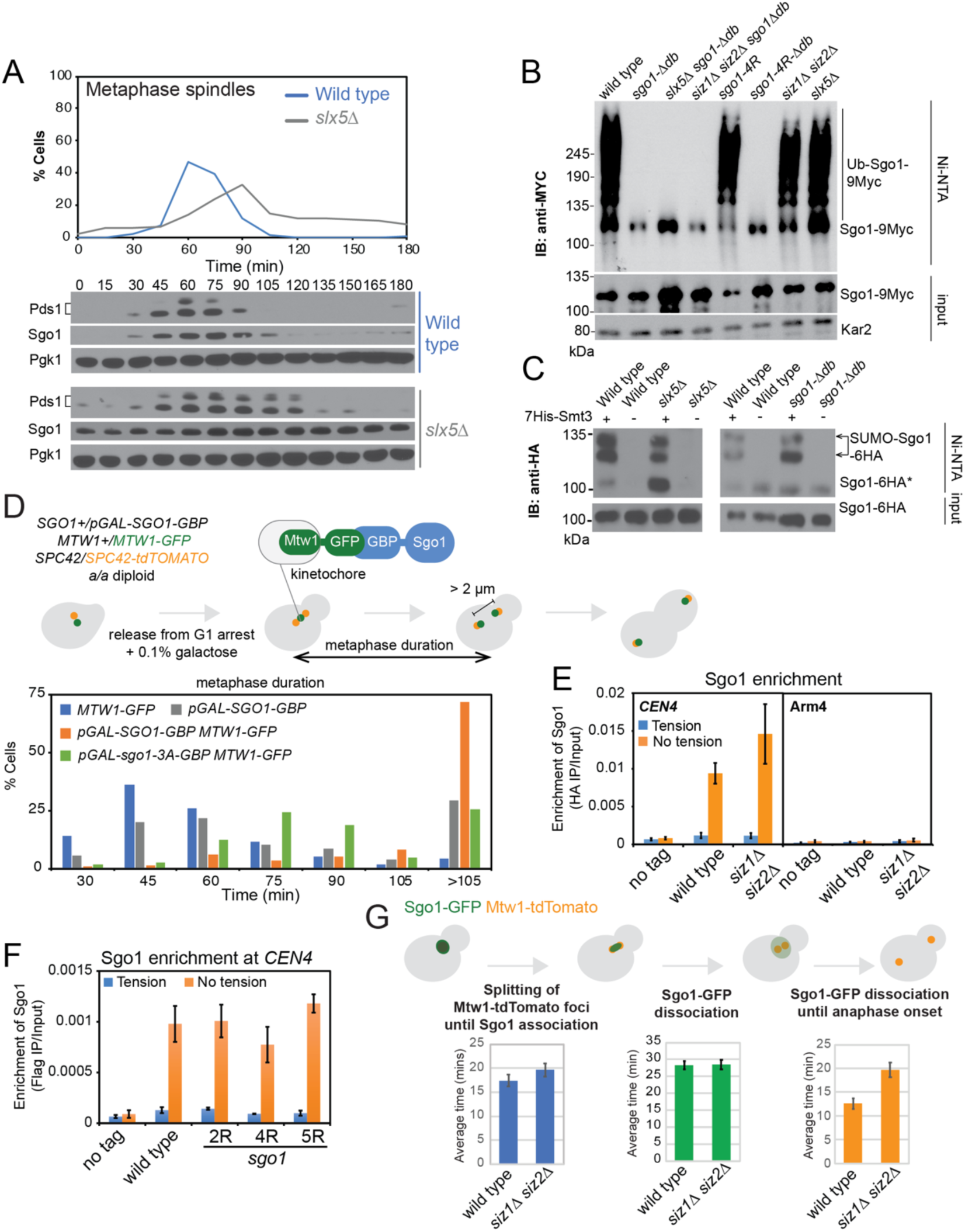
SUMOylation promotes anaphase onset independently of Sgo1 degradation or removal from pericentromeres. (A) *slx5*Δ cells exhibit a metaphase delay and stabilize Sgo1. Wild type (AMy8467) and *slx5*Δ (AMy10981) strains carrying *PDS1-3HA* and *SGO1-9MYC* were analysed as described in Figure 1F. (B) Sgo1 ubiquitination is dependent on its destruction box and independent of Sgo1 SUMOylation or Slx5/Slx8. Strains with *pGAL*-*SGO1* (AMy27029), *pGAL*-*sgo1-Δdb* (AMy27030), *slx5*Δ *pGAL*-*sgo1-Δdb* (AMy27031), *siz1*Δ *siz2*Δ *pGAL*-*sgo1-Δdb* (AMy27032), *pGAL-sgo1-4R* (AMy27033), *pGAL*-*sgo1-4R-Δdb* (AMy27034), *siz1*Δ *siz2*Δ *pGAL*-*SGO1* (AMy27035) and *slx5*Δ *pGAL*-*SGO1* (AMy27036) and carrying His-UBI (AMp1673) were arrested in G1 in raffinose medium and *pGAL*-*SGO1-9MYC* expression was induced by the addition of galactose. Ubiquitinated proteins were purified on Ni-NTA resin and Sgo1-9Myc was detected in inputs and elutes by anti-Myc immunoblot. (C) Sgo1*Δdb* and *slx5*Δ do not noticeably change the levels of Sgo1-SUMO. *In vivo* SUMO assay was performed as described in Figure 2A. Cells analysed were wild type (AMy906), *slx5*Δ (AMy9765) and *sgo1-Δdb* (AMy18153), carrying Sgo1-6HA. (D-G) Sgo1 dissociation from the pericentromeres is important for promoting anaphase onset, but occurs independently of SUMOylation. (D and E) Tethering Sgo1 to the kinetochore component Mtw1 blocks anaphase onset. (D) Scheme of the experiment. Live cell imaging was performed in a/a diploid synchronised by release from G1. Metaphase duration was determined as the time between the observation of two Spc42-tdTomato dots until they reached a distance of > 2 μm apart. (E) Metaphase duration was measured in at least 200 *pGAL-SGO1-GBP* (AMy26679), *MTW1-GFP* (AMy26682), *pGAL-SGO1-GBP MTW1-GFP* (AMy26568) and *pGAL-sgo1-3A-GBP MTW1-GFP* (AMy26570) cells carrying *SPC42-tdTOMATO*. (F) SUMOylation is not required for Sgo1 dissociation from the pericentromeres under tension. No tag control (AMy2508), *SGO1-6HA* (AMy6390) and *siz1*Δ *siz2*Δ *SGO1-6HA* (AMy8115) cells carrying *MET-CDC20* were arrested in metaphase in the presence (DMSO) or absence (benomyl/nocodazole) of spindle tension and Sgo1 association with the indicated site was measured by ChIP-qPCR. (G) SUMO-deficient Sgo1 mutants show similar localization to wild type Sgo1. Sgo1 association with *CEN4* was measured by ChIP-qPCR using wild type (AMy25141), *sgo1-2R* (AMy26707), *sgo1-4R* (AMy26696) and *sgo1-5R* (AMy26708) strains carrying *SGO1-6HIS-3FLAG*, together with a no tag control (AMy2508). Cells were arrested in metaphase by depletion of Cdc20 in the presence or absence of spindle tension. (H) *siz1*Δ *siz2*Δ delay in metaphase after bulk Sgo1 removal from the pericentromere. Wild type (AMy9233) and *siz1*Δ *siz2*Δ (AMy15604) strains carrying *MET-CDC20*, *SGO1-yeGFP* and *MTW1-dtTOMATO* were followed by live cell imaging and the time elapsed between the indicated events was quantified for at least 50 cells. Anaphase onset was estimated as the time when the two Mtw1-tdTOMATO dots were > 2 μm apart.

To examine whether Sgo1 could be a direct target of Slx5/Slx8 we analysed ubiquitin conjugates in *siz1*Δ *siz2*Δ, *slx5*Δ and *sgo1-*4R cells. Since Sgo1 degradation also requires APC/C-dependent ubiquitination within a C-terminal degradation box on Sgo1 [14], we also analyzed mutants lacking this region (*Δdb*). Strikingly, Sgo1 ubiquitination was abolished in cells lacking the APC/C-dependent degradation box, but not detectably reduced in *siz1*Δ *siz2*Δ, *slx5*Δ or *sgo1-*4R cells, indicating that Sgo1 is unlikely to be a direct target of STUbL enzymes (Figure 4B). Consistently, Sgo1-SUMO conjugates do not accumulate in *slx5*Δ cells (Figure 4C) and the half-life of Sgo1-2R was not increased over wild type Sgo1 (Figure S4D). Therefore, Slx5/Slx8 regulates anaphase onset, but not by directly ubiquitinating Sgo1, and although Siz1/Siz2 and Slx5 affect Sgo1 stability, they do so indirectly, for example, by activating proteosome function [24].

Moreover, Sgo1 degradation appears dispensable for anaphase onset since *sgo1*-Δ*db* cells show no metaphase delay [14] and abolishing the APC/C recognition sites in Sgo1 did not exacerbate the metaphase delay of the *sgo1-2R* mutant (*sgo1-2R* Δ*db*, Figure S3D and S4E). Importantly, Sgo1-Δdb was SUMOylated to a similar extent to wild type Sgo1 (Figure 4C). Therefore, Sgo1 SUMOylation is able to promote anaphase onset even in the absence of APC/C-mediated degradation.

### Sumoylation does not promote Sgo1 removal from chromatin under tension

Sgo1 is released from pericentromeres under tension [8] but whether this is critical for the metaphase-anaphase transition remained unclear. To address this, we asked if artificial tethering of Sgo1 to the kinetochore can prevent tension-dependent removal: GFP-binding protein (GBP) -tagged Sgo1 was produced from a galactose-inducible promoter (replacing endogenous Sgo1) in cells where the kinetochore protein Mtw1 was tagged with GFP (Mtw1-GFP) (Figure 4D). On its own, *pGAL*-*SGO1-GBP* expression caused a modest metaphase delay compared to the Mtw1-GFP control, presumably due to mild overexpression (Figure 4D). However, Sgo1-GBP expression in a strain producing Mtw1-GFP resulted in a severe delay in metaphase (Figure 4D). This delay required Sgo1 association with PP2A-Rts1 and/or CPC, because kinetochore tethering of the Sgo1-3A mutant protein, which has lost these interactions, resulted in a more modest metaphase delay (Figure 4D). Therefore, kinetochore-associated Sgo1 prevents anaphase onset in a manner depending on its ability to bind PP2A-Rts1 and/or CPC, showing that tension-dependent removal of Sgo1 is critical for anaphase entry.

Based on these findings, we considered that Siz1/Siz2 may promote anaphase entry by triggering the release of Sgo1 from chromosomes upon sister kinetochore biorientation. In wild type metaphase-arrested cells, chromatin immunoprecipitation followed by qPCR (ChIP-qPCR) showed that Sgo1 associates with a centromeric site in the absence, but not presence, of spindle tension and this pattern was largely unchanged in *siz1*Δ *siz2*Δ, *sgo1-2R, sgo1-4R* or *sgo1-5R* cells with reduced SUMOylation (Figure 4E and F; Figure S4F and G). We confirmed that SUMOylation is dispensable for the tension-dependent release of Sgo1 during anaphase by live cell imaging. In wild type cells, Sgo1-GFP first appeared as a bright focus which dissociated upon splitting of Mtw1-tdTomato foci at metaphase and this occurred with similar timing in *siz1*Δ *siz2*Δ cells (Figure 4G, Figure S4H). Instead, the observed delay in anaphase entry occurred after bulk Sgo1 removal (Figure 4G). Hence, although tension-dependent release of Sgo1 is critical for anaphase entry, this occurs independently of Sgo1 SUMOylation.

### Sgo1 SUMOylation is required for stabilizing biorientation

Our findings indicate that SUMOylation neither targets Sgo1 for STUbL-mediated destruction nor does it facilitate Sgo1 removal under tension. To probe the mechanism connecting Sgo1 SUMOylation to timely anaphase onset, we visualized the efficiency of sister kinetochore biorientation in the Sgo1 SUMO mutants. We analysed the initial establishment of sister kinetochore biorientation by monitoring the separation of sister *CEN4-GFP* foci as spindles reformed after nocodazole wash-out, while maintaining a metaphase arrest (by depletion of Cdc20, Figure 5A). In contrast to *sgo1-3A* cells which have impaired biorientation, likely due to impaired CPC binding[10], all mutants with reduced Sgo1 SUMOylation except *sgo1-5R* (*siz1*Δ *siz2*Δ, *sgo1-2R* and *sgo1-4R*) showed proficient sister kinetochore biorientation (Figure 5B). Therefore, error correction and sister kinetochore biorientation pathways are functional in the absence of Sgo1 SUMOylation.

Next, we assessed the stability of biorientation in the SUMO mutants. Cells were released from nocodazole washout, and the separation of *CEN4-GFP* was monitored as cells progressed into anaphase. A single *CEN4-GFP* focus was observed initially, and two *CEN4-GFP* foci appeared upon attachment of sister kinetochores to microtubules from opposite poles. Stable attachment led to further separation of the two *CEN4-GFP* foci, which eventually segregated to opposite poles in anaphase (Figure 5C). In contrast, if attachments are unstable, the two *CEN4-*GFP foci reassociate prior to their splitting and segregation. In *sgo1*Δ and *sgo1-3A* mutants, which are defective in sensing and correcting attachment errors, the visualisation of two *CEN4-GFP* foci was delayed and the number of missegregation events was increased, but the levels of reassociation of two *CEN4-GFP* foci was similar to wild type (Figure 5D and E, Figure S5A and B). *siz1*Δ *siz2*Δ and *sgo1-4R* mutants, in stark contrast, were proficient in the initial establishment of biorientation and did not show increased missegregation (Figure 5D and E, Figure S5A and B). Instead, both mutants showed ∼15% increase in the number of cells in which the two *CEN4-GFP* foci reassociated, indicative of unstable biorientation. We conclude that Sgo1 SUMOylation is important to maintain the bioriented state.

### The metaphase delay in siz1Δ siz2Δ is rescued by inactivating mutations in CPC/SAC

Unstable biorientation in SUMO-deficient mutants is expected to generate unattached kinetochores and engage the SAC, potentially explaining the metaphase delay of these cells. Consistent with this idea, deletion of *MAD2* partially rescued the metaphase delay of *siz1*Δ *siz2*Δ cells (Figure S6A). The CPC-dependent error correction pathway is likely responsible for the instability of kinetochore-microtubule interactons in the SUMO mutants because inhibition of the CPC component, Ipl1 (*ipl1-as1*) also reduced the metaphase delay of *siz1*Δ *siz2*Δ cells (Figure 6A). Consistently, Sgo1 interaction with PP2A-Rts1 and/or CPC, is important for the metaphase delay in the absence of SUMOylation because both time course analysis (Figure S6B) and live cell imaging (Figure 6B) revealed that *sgo1-3A siz1*Δ *siz2*Δ cells spent less time in metaphase than *siz1*Δ *siz2*Δ cells. Interestingly, however, unlike Ipl1 (Figure 6A), the PP2A regulatory subunit, Rts1, was largely dispensible for the metaphase delay of *siz1*Δ *siz2*Δ cells (Figure S6C). Therefore, the CPC-dependent error correction pathway is responsible for the metaphase delay observed in the absence of Sgo1 SUMOylation.

**Figure 6.**
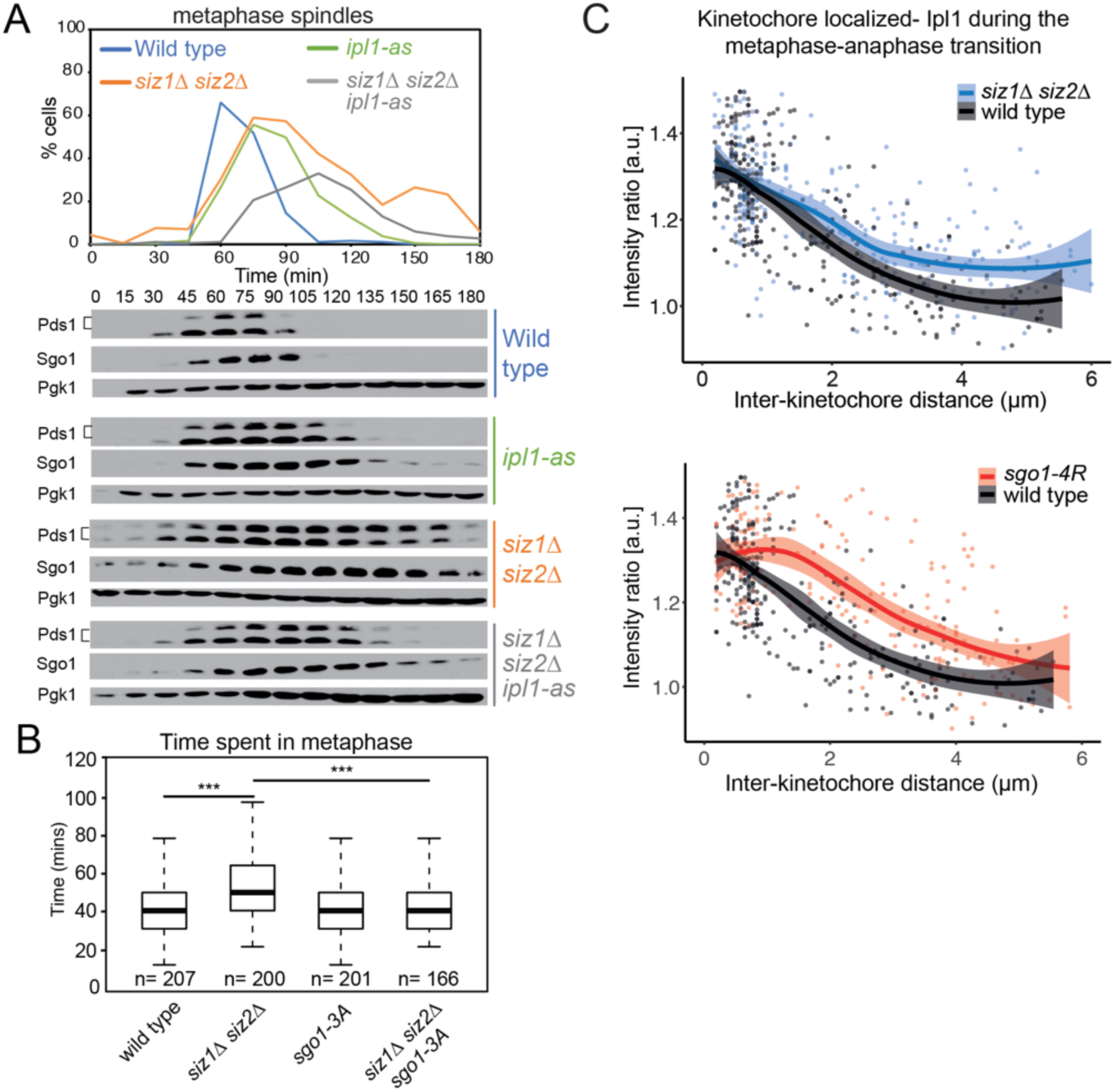
Sgo1 SUMOylation promotes timely removal of Ipl1 from kinetochores. (A) Inhibition of Ipl1 kinase activity partially rescues the metaphase delay phenotype of *siz1*Δ *siz2*Δ mutant. Cells were released from G1 arrest, NA-PP1 was added when small buds emerged, and cell cycle progression was assessed as described in Figure 1F. Strains used were wild type (AMy8467), *ipl1-as5* (AMy15026), *siz1*Δ *siz2*Δ (AMy8452) and *siz1*Δ *siz2*Δ *ipl1-as5* (AMy15237). (B) Metaphase duration was determined by live cell imaging as described in Figure 1D. Strains used carried *TUB1-YFP* and *CDC14-GFP* and were wild type (AMy24174), *siz1*Δ *siz2*Δ (AMy24313), *sgo1-3A* (AMy24433) and *siz1*Δ *siz2*Δ *sgo1-3A* (AMy24471). n = number of cells analyzed. Data were presented as box plots, with the median values indicated by the thick black lines, upper and lower quartiles marked by the upper and lower borders of the boxes, and maximum (excluding outliers) and minimum values marked by the error bars. *P-*values were calculated using the Mann-Whitney U-test. *** = *P* < 0.001. (C) Ipl1 stays longer on kinetochores during the metaphase-anaphase transition in Sgo1 SUMO-deficient mutants. Strains used contained *IPL1-yeGFP*, *MET-CDC20* and *MTW1-tdTOMATO*, and were wild type (AMy9231), *siz1*Δ *siz2*Δ (AMy15602) and *sgo1-4R* (AMy24143). Cells were released from G1 and were imaged on a microfluidics device. Line scans were performed across kinetochore foci of single cells, which measured the distance between the two Mtw1-tdTOMATO foci, as well as the Ipl1-GFP intensities co-localizing with the Mtw1 foci. Intensity ratio = the average intensity of the two Ipl1-GFP signals/average intensity of the two Mtw1-tdTOMATO signals. Shaded areas represent 95% confidence intervals. > 200 line scans were performed for each strain.

### Sgo1 SUMOylation promotes Ipl1 relocalization

The kinase activity of Ipl1 (Aurora B kinase) is required for error correction and Ipl1 is known to re-localize from centromeres to the spindle mid-zone upon the establishment of biorientation [25, 26]. Unstable biorientation in Sgo1 SUMOylation mutants suggested that this removal may be incomplete. We monitored Ipl1-GFP and its co-localization with Mtw1-tdTomato by imaging cells released from G1. In both *siz1*Δ *siz2*Δ and *sgo1-*4R, Ipl1 recruitment to the kinetochore-proximal regions occurred normally (Figure 6C). However, as kinetochores separated, Ipl1-GFP persisted close to kinetochores in the Sgo1 SUMO mutants (Figure 6C). Increased centromeric Ipl1 was also measured by ChIP in SUMO-deficient cells arrested in metaphase with kinetochores under tension (Figure S7A and B). Hence, Sgo1 SUMOylation facilitates the re-localization of Ipl1from centromeres to prevent persistent error correction and SAC activation.

### Sgo1 SUMOylation is incompatible with PP2A-Rts1 binding

Interestingly, the coiled-coil domain of Sgo1 is both required for its SUMOylation (Figure 3) and for PP2A-Rts1 binding [10, 27]. This raised the question of whether Sgo1 SUMOylation also impacts PP2A-Rts1 binding. Sgo1 SUMOylation was increased in the Sgo1-3A mutant (Figure 7A), suggesting that PP2A-Rts1 binding normally dampens Sgo1 SUMOylation. Structural modelling of budding yeast Sgo1-PP2A-Rts1 interaction revealed that PP2A-Rts1 binding to Sgo1 would be incompatible with SUMOylation on these residues (Figure S7C). We used an *in vitro* binding assay to test the effects of Sgo1 SUMOylation on Rts1 binding. Purified Sgo1 was SUMOylated on beads *in vitro*, beads were stringently washed to remove components of the SUMO reaction and subsequently incubated with cell-free extract from *sgo1*Δ Rts1-9Myc cells. This revealed that, as expected, Rts1-9Myc bound robustly to unSUMOylated Sgo1, however Rts1 binding was greatly reduced by Sgo1 SUMOylation, consistent with the prediction that SUMOylation and PP2A-Rts1 binding are mutually exclusive (Figure 7B). Similarly, immunoprecipitated Rts1-9Myc bound *in vitro* SUMOylated Sgo1 less well than unSUMOylated Sgo1 (Figure 7C). Analysis of the SUMO-deficient Sgo1-4R protein showed that, unexpectedly, binding of Rts1 was reduced to a similar extent to Sgo1-3A, even in the absence of SUMOylation (Figure 7B). Despite the fact that both mutants fail to bind PP2A-Rts1, they result in very different outcomes *in vivo*: while *sgo1-3A* shows defective initial biorientation of sister kinetochores and chromosome mis-segregation after nocodazole washout, *sgo1-4R* does not (Figure 5B and E). Conversely, unstable sister kinetochore biorientation and a metaphase delay is observed in *sgo1-4R* but not *sgo1-3A* cells (Figure 3D, Figure 5D, Figure 6B, Figure S6B). This indicates that a failure to bind PP2A-Rts1 cannot be the primary cause of these defects. Instead, sister kinetochore biorientation and chromosome segregation defects in *sgo1-3A* cells are attributed to defective CPC maintenance at kinetochores [10] while, conversely, our data indicate that CPC persists at kinetochores in *sgo1-4R* cells causing unstable biorientation and a metaphase delay (Figure 6C). Collectively, these data indicate that the ability of Sgo1 to bind and release CPC underlies the establishment and stabilization of biorientation, respectively (Figure S7D).

**Figure 7.**
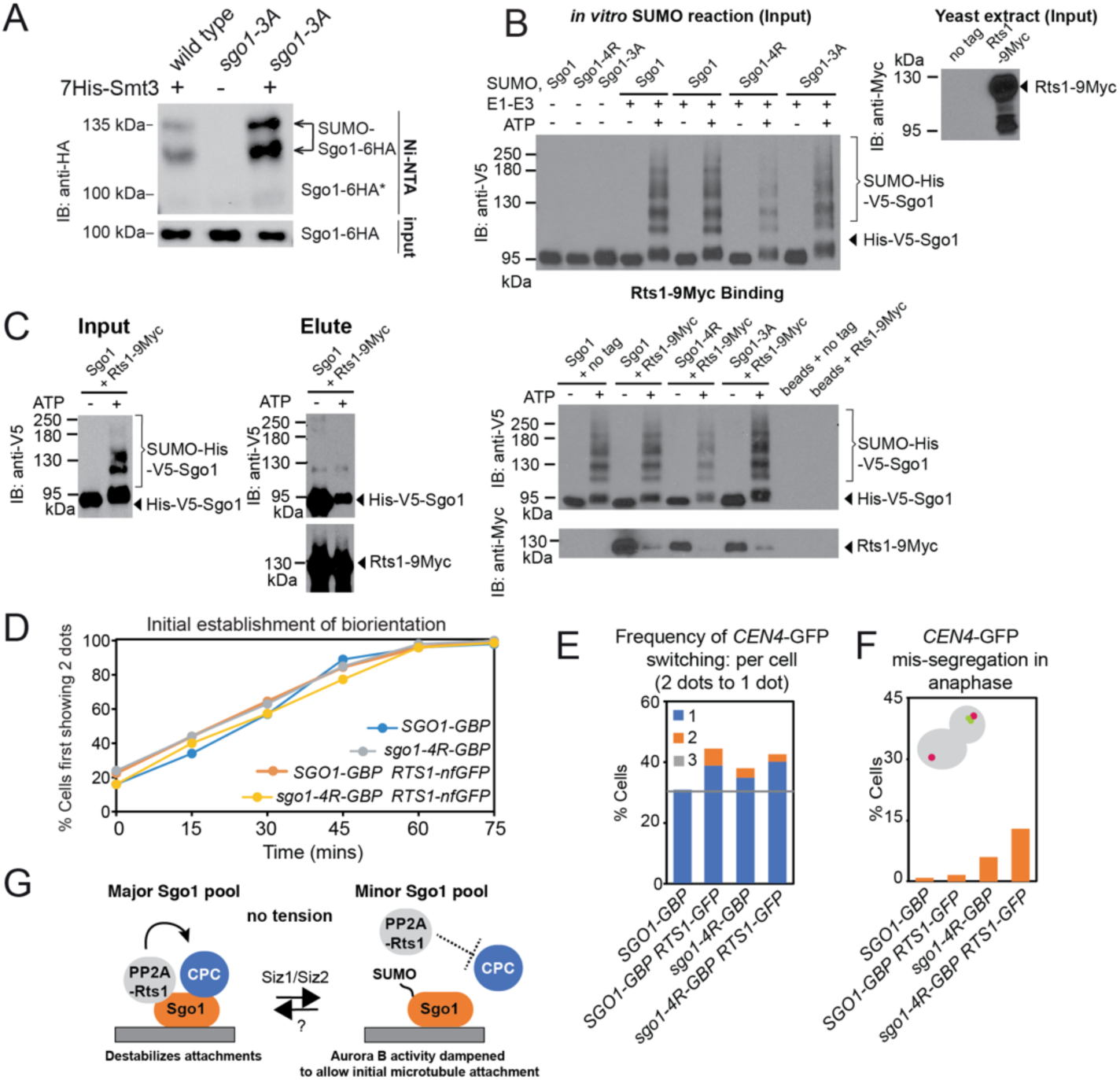
SUMOylation blocks Rts1 binding to Sgo1 and release of this interaction is important for stable biorientation. (A) The Sgo1-3A mutant which fails to associate with PP2A-Rts1 or CPC shows enhanced SUMOylation. *In vivo* SUMOylation assay was performed on wild type (AMy7655) and *sgo1-3A* (AMy25988) strains carrying *SGO1-6HA*. (B) SUMOylated Sgo1 has reduced binding affinity for Rts1. Recombinant V5-tagged Sgo1 was mixed with components of the SUMOylation pathway in the presence or absence of ATP. Anti-V5 antibody coupled Protein G dynabeads were added to the mixture, washed thoroughly and incubated with extract from *sgo1*Δ or *sgo1*Δ *RTS1-9MYC* (AMy8832). Levels of Sgo1 and Rts1 bound to beads were probed by anti-V5 and anti-Myc western blotting, respectively. (C) Rts1 preferentially binds to unsumoylated Sgo1. Rts1-9MYC was immunoprecipitated from *sgo1*Δ *RTS1-9MYC* (AMy8832) using anti-MYC antibody coupled to Protein G dynabeads. Beads were incubated with *in vitro* SUMOylation reaction mixture containing Sgo1. Levels of Sgo1 and Rts1 bound to beads were probed by anti-V5 and anti-Myc western blotting, respectively. (D-F) Biorientation is unstable when Rts1 is tethered to wild type Sgo1 or Sgo1-4R. Strains used contained *CEN4-mNeonGreen MET-CDC20* and *SPC42-tdTOMATO* and were *SGO1-GBP* (AMy28389), *SGO1-GBP RTS1-non-fluorescent GFP* (AMy28092), *sgo1-4R-GBP* (AMy28417) and *sgo1-4R-GBP RTS1-non-fluorescent GFP* (AMy28416). The assay was performed as described in Figure 5C. (D) Tethering Rts1 to wild type Sgo1 or Sgo1-4R does not affect the initial establishment of biorientation. (E) Increased reassociation of *CEN4-mNeonGreen* dots was observed when Rts1 was tethered to wild type Sgo1 or Sgo1-4R. (F) Mis-segregation is only modestly increased when Rts1 is tethered to wild type Sgo1 or Sgo1-4R. (G) Schematic model of how Sgo1 SUMOylation may alter the kinase-phosphatase balance to initiate error correction silencing and promote anaphase onset. For details, see text

### Dissociation of shugoshin and PP2A-Rts1 stabilizes sister kinetochore biorientation

To understand the importance of the Sgo1-PP2A-Rts1 interaction in stabilizing sister kinetochore biorientation, we asked whether tethering of Rts1 to wild type Sgo1 or Sgo1-4R could rescue the instability of attachments. Although forcing Rts1 interaction with Sgo1-4R throughout the cell cycle did not affect initial biorientation (Figure 7D), this state was unstable, as judged by the increased number of cells switching between two and one *CEN4-GFP* dots (Figure 7E), though chromosome segregation was ultimately successful in the majority of cells (Figure 7F). This further confirms that loss of Rts1 binding is not the cause of unstable sister kinetochore biorientation in *sgo1-4R* cells. However, interestingly, forced interaction between Rts1 and wild type Sgo1 also resulted in frequent switches (Figure 7E). Therefore, release of the Sgo1-Rts1 interaction is important to stabilize bioriented sister kinetochore-microtubule attachments. Together with our finding that PP2A-Rts1 binding is incompatible with Sgo1 SUMOylation, this suggests that PP2A-Rts1 dissociation as a result of Sgo1 SUMOylation, is also important to stabilize biorientation.

## Discussion

### Identification of shugoshin regulators

Starting with an unbiased genetic screen we have identified SUMO ligases as negative regulators of the pericentromeric hub that responds to a lack of tension between kinetochores. Sgo1, the central pericentromeric adaptor protein is one key target of the Siz1/Siz2 SUMO ligases. Kinetochore-microtubule interactions are unstable in Sgo1-SUMO deficient cells (*siz1*Δ *siz2*Δ and *sgo1-4R*). Persistent cycles of kinetochore detachment and re-attachment engage the SAC, explaining why a failure to SUMOylate Sgo1 results in a metaphase delay. Consistently, we find that inactivation of components of the error correction pathway (Ipl1, Mad2) advanced anaphase timing in *siz1*Δ *siz2*Δ cells.

Sgo1 inactivation and removal from kinetochores is essential for timely anaphase entry ([18], Figure 4D) and here we have identified one mechanism that contributes to this inactivation. However, Sgo1 also prevents anaphase onset by inhibiting separase independently of securin [18]. PP2A-Cdc55 dependent dephosphorylation of separase and, potentially also cohesin itself is likely to underlie this inhibition [28, 29]. Notably, *ZDS2*, a negative regulator of PP2A-Cdc55 [30], was also isolated in our screen along with *HOS3*, the cohesin deacetylase [31–33] indicating that further mechanisms await discovery.

### SUMOylation of Sgo1 ensures efficient entry into anaphase

How does Sgo1 SUMOylation regulate anaphase entry? We found that Sgo1 SUMOylation is dispensable for its tension-dependent release from pericentromeres and that, although SUMOylation promotes Sgo1 degradation indirectly, this is not required for efficient anaphase entry. Instead, our work suggested that Sgo1 SUMOylation likely promotes anaphase entry by silencing the error correction process, as biorientation was highly unstable in Sgo1 SUMO-deficient mutants (Figure 5D). Remarkably, we found that Ipl1 removal from kinetochores was incomplete in the Sgo1 SUMO mutants (Figure 6C), suggesting a key role of this modification in modulating the subcellular localization of Ipl1.

Meanwhile, we showed that Rts1 binds preferentially to unSUMOylated Sgo1, and that tethering Sgo1 to Rts1 destabilized biorientation in a similar way as the Sgo1 SUMO mutants. These findings suggest that Sgo1 SUMOylation-mediated Rts1 dissociation has an important role in stabilizing microtubule-kinetochore attachment. Interestingly, PP2A-B56 dampens the effects of Aurora B to allow initial attachments in human cells [34–36], suggesting a potential explanation for our observations. Overall, our findings indicate that SUMOylation modulates the kinase-phosphatase balance at the kinetochore to dampen CPC activity and allow initial kinetochore-microtubule attachments (Figure 7G). Different mutants affect this balance in distinct ways, leading to the observed outcomes on the establishment and stabilization of biorientation (Figure S7D).

### SUMOylation – a generalized mechanism of centromere regulation with implications for disease?

Accumulating evidence indicates that SUMOylation might play a specific role at centromeres to fine-tune chromosome segregation. The SUMO isopeptidase Ulp2/Smt4 is important for maintenance of cohesion specifically at centromeres, in part through regulating Topoisomerase II [37]. The Pds5 subunit of cohesin is also known to prevent polySUMOylation of cohesin [38] and centromeric cohesin may be particularly susceptible since it lacks Pds5 [39]. PIAS SUMO ligases are known to be localized at centromeres in vertebrate mitotic cells and oocytes [40–42]. Moreover, the SUMO pathway is required to prevent cohesion loss during meiosis II, which centrally requires Sgo2-PP2A, and it is conceivable that modulation of the PP2A-Sgo1 interaction as we find in yeast underlies these observations in mouse oocytes [43]. Indeed, global studies found that shugoshins are SUMOylated in fission yeast and human cells, though the function has yet to be tested [44, 45]. Furthermore, there is ample evidence that CPC function may be subject to regulation by SUMOylation. Aurora B SUMOylation in *Xenopus* and human cells was shown to promote its enrichment at centromeres and it was proposed that this modification may serve as a reversible mechanism to dampen Aurora B kinase activity [46], while in *C. elegans* meiosis the localization of both Aurora B and Bub1 kinase are influenced by SUMOylation [47–49]. This suggests that SUMO may have a general role in CPC/error correction pathways and we speculate that multi-lateral SUMO-SIM interactions [50] enable the coordinated re-localization of surveillance factors. Notably, mis-regulation of the SUMO pathway is wide-spread in different cancers [51, 52]. Potentially, reductions in chromosome segregation fidelity caused by SUMO malfunction at centromeres, as we show here, could be a contributing factor in driving these malignant states.

## Acknowledgments

We are grateful to Yoshiko Kikuchi and Helle Ulrich for yeast strains and plasmids, respectively, to Weronika Borek for help with R programming. We thank Lorraine Pillus, Emily Petty, Federico Pelisch and Andrew Goryachev for helpful discussions and comments on the manuscript. This work was funded by Wellcome through Senior Research Fellowships to AM [[107827]] and AAJ [202811], and core funding for the Wellcome Centre for Cell Biology [203149].

## Author contributions

Conceptualization, A.L.M., X.B.S., C.S., L.W., O.N., D.C., A.W. and Z.S.; Methodology, X.B.S, M.W., C.S., O.N., T.T. and R.H. Investigation: X.B.S., M.W., C.S., D.C., O.N., A.W., T.T. and Y.W. Software: D.A.K. Writing – original draft, X.B.S.. and A.L.M.; Writing – Review and Editing, all authors; Visualization, X.B.S., A.A.J. and A.L.M.; Supervision, A.L.M., R.H. and Z.S.; Funding acquisition, A.L.M.

## Declaration of interests

The authors declare no competing interests.

## Methods

### Yeast strains and plasmids

Yeast strains are derivatives of W303 and are listed in Supplementary Table S3. Plasmids and primers are listed in Supplementary Tables S4 and S5, respectively. StuI-digested AMp1239 was transformed into a *CDC14-GFP* strain to make the *YFP-TUB1 CDC14-GFP* parent strain. Genes were deleted or tagged using PCR-based transformation. K-R mutant plasmids were generated using Quikchange II XL site-directed mutagenesis kit (Agilent), with primers listed in Table S5. K-R mutants were PCR amplified from the resulting plasmid using primers AM16 and AM3177 and were integrated into an *sgo1*Δ strain (AMy827). *7HIS-SMT3* and *HIS-UBI* plasmids were kind gifts from Dr. H. Ulrich.

### Yeast growth and synchronization

Unless otherwise stated, yeast strains were grown in YEP supplemented with 2% glucose and 0.3 mM adenine (YPDA). For the benomyl sensitivity assays, plates were made by adding 10 μg/mL benomyl (Sigma) or the equivalent volume of DMSO (solvent control) to boiling media.

To synchronize cells in G1, overnight cultures were inoculated to OD_600_ = 0.2 and grown for 1 h at room temperature, before diluting back to OD_600_ = 0.2. α-factor was added to 5 μg/mL for 90 min and then re-added to 2.5 μg/mL for every 90 min, until > 95% cells exhibited shmooing morphology. To release cells from G1, α-factor was washed out using 10 × volume relevant media. For *pMET-CDC20*-containing strains, cells were arrested in G1 in methionine dropout medium. After α-factor wash-out, cells were released into YPDA (+ DMSO or + 30 μg/mL benomyl and 15 μg/mL nocodazole) + 8 mM methionine for 1 h. 4 mM methionine and DMSO/15 μg/mL nocodazole were re-added for 1 h.

### Metaphase duration measurements by live cell imaging and mitotic time course

Synthetic complete/dropout media were used for growing and washing cells for live cell imaging. Cells released from G1-arrest were loaded onto μ-slide dishes (Ibidi) coated with concanavalin-A (Sigma). Images were taken at indicated time intervals using a Zeiss inverted microscope, in a temperature-controlled chamber (25°C for glucose-based media and 30°C for raffinose-based media). Movies were analyzed using the ImageJ software.

Mitotic time course experiments were performed at 25°C. To inhibit Ipl1-as5, NA-PP1 (Toronto Research Chemicals) was added to a final concentration of 50 μM upon the emergence of small-budded cells. For every time point, samples were either treated with 5% trichloroacetic acid (TCA) for protein extraction, or fixed in 3.7% formaldehyde for immunofluorescence. TCA-treated pellets were snap frozen in liquid nitrogen, washed in acetone and air-dried. Protein samples were prepared by bead beating and boiling in SDS sample buffer. For immunofluorescence, Fixed cells were spheroplasted using zymolyase (AMS Biotechnology) and glusulase (Perkin Elmer), fixed in methanol for 3 min, followed by 10 s incubation in acetone. Rat anti-α-tubulin (Abd Serotec) antibody was used at 1:50 and anti-Rat FITC (Jackson ImmunoResearch) was used at 1:16.7. Spindle morphology of 200 cells was analyzed for each time point.

### Western blotting

Proteins were separated in 8% bis-tris acrylamide gels and were transferred to nitrocellulose membranes (except anti-ubiquitination blots, for which PVDF membranes were used). Membranes were blocked in 3-5% milk in phosphate-buffered saline + 0.1% Tween-20 (PBST) or tris-buffered saline + 0.1% Tween-20 (TBST). All antibodies were diluted in 2% milk PBST (except anti-ubiquitination blot, for which 2% milk TBST was used). Anti-c-Myc (9E10, Biolegend), anti-V5 (AbD Serotec), anti-Flag (M2, Sigma) and anti-HA (12CA5, Roche) antibodies were used at 1:1000 dilution. Anti-Pgk1 (lab stock) and anti-Kar2 (lab stock) loading controls were used at 1:10000 and 1:100,000 dilution, respectively. The non-quantitative blots were detected by ECL (Thermofisher) and autoradiograms. 20% Femto-ECL (Thermofisher) diluted in Pico-ECL was used to detect SUMOylated-Sgo1 signals. Quantitative blots were detected using a BioRad Chemidoc (Figure S4C) or a LiCOR Odyssey Clx machine (Figure S4D). Protein expression was quantified using ImageJ (Figure S4C) or ImageStudio (Figure S4D).

### Analysis of *in vivo* SUMOylation

Cultures were inoculated to OD_600_ = 0.2 in 200 mL synthetic dropout media and grown for 4 h at room temperature. Equal OD of cells were collected for samples in the same experiment. Cell pellets were resuspended in 20 mL cold H_2_O and incubated with 3.2 mL solution containing 1.85 M sodium hydroxide and 7.5% β-mercaptoethanol. After 20 min incubation on ice, 1.65 mL 100% trichloroacetic acid was added and cells were incubated on ice for a further 20 min. Cell pellets were drop-frozen in liquid nitrogen, and subsequently lysed by bead-beating in 300 µL buffer A (6 M guanidine hydrochloride, 100 mM sodium phosphate buffer pH 8.0, 10 mM Tris-HCl pH 8.0). Lysate was diluted three-fold in buffer A and 10 µL was saved as input controls. Lysate was applied to a column packed with 600 μL 50% slurry Ni-NTA agarose beads (Qiagen), washed twice with buffer A + 0.05% Tween-20, twice with buffer B (8 M urea, 100 mM sodium phosphate buffer pH 6.8, 10 mM Tris-HCl pH 6.8) + 0.05% Tween-20 and once with buffer B + 0.05% Tween-20 + 20 mM imidazole. SUMOylated proteins were eluted by buffer B + 0.05% Tween-20 + 200 mM imidazole. Input and elute samples were concentrated by centrifugation using Vivaspin columns (Sartorius).

### Mass spectrometry

A *pGAL-SGO1* strain with *7HIS-SMT3* was inoculated to 0.2 OD in 2% raffinose media. After 3 h growth at room temperature, cells were diluted back to 0.2 OD and 2% galactose was added to induce *SGO1* overexpression. Cells were harvested after 3 h and SUMOylated proteins were purified as described above. Proteins eluted from the Ni-NTA column were separated on a 4-12% NuPage Bis-Tris gel (Invitrogen) and the gel slice encompassing SUMOylated Sgo1 (between ∼100 kDa and 135 kDa, based on immunoblotting of a parallel gel) was excised for mass spectrometry analysis.

### Cloning, expression and purification of recombinant Sgo1

Full-length wild type *SGO1* was amplified from plasmid AMp899, to replace *SMT3* in plasmid AMp773 by Gibson assembly using primers AM8849 to AM8852. V5 tag was inserted by Gibson assembly using plasmid AMp970 and primers AM8866 to AM8869 to generate N-terminal tobacco etch virus (TEV) protease cleavable His_x7_-V5 tag-tagged *SGO1* (AMp1738) under the control of a *pCUP1* promoter. *sgo1-4R* was PCR amplified from plasmid AMp1340 and ligated into AMp1738 by Gibson assembly using primers AM8850, AM8851, AM9124 and AM 9125 to generate N-terminal TEV protease cleavable His_x7_-V5 tag-tagged *sgo1-4R* (AMp1783).

A protease-deficient yeast strain (AMy8184) was transformed with the resulting plasmids (AMp1738 or AMp1783) and the transformants were inoculated into 8 L uracil dropout media. When OD_600_ reached 0.5 - 0.7, 0.5 mM CuSO_4_ was added to induce His_x7_-V5 -*SGO1* (or *sgo1-4R*) expression. Cells were harvested 6 h after induction. Cell pellets were snap frozen in liquid nitrogen and ground to powder in a ball breaker machine (Retsch). All purification steps were performed at 4°C or on ice. Cell powder was resuspended in lysis buffer (25 mM Tris-HCl pH 7.5, 150 mM NaCl, 1 mM MgCl_2_, 10 % glycerol, 0.1% NP-40, 0.05 mM EDTA, 0.05 mM EGTA, 1 mM DTT, 1xCLAAPE, 1 mM Pefabloc, 0.4 mM Na Orthovanadate, 0.1 mM Microcystin, 1 mM NEM, 2 mM B-glycerophosphate, 1 mM Na pyrophosphate, 5 mM NaF, complete EDTA-free protease inhibitor (Roche)). Cell lysates were treated with 40 u/mL benzonase for 1.5 h. The crude lysate was diluted with 25 mM sodium phosphate buffer pH 7.5, 500 mM NaCl, 10 % glycerol, 0.1% NP-40, 10 mM imidazole, 1 mM DTT, 0.25 mM PMSF, cleared by ultracentrifugation (50,000 ×g at 4°C for 30 min), filtered through a 0.22 µm filter and loaded onto a HiTrap IMAC FF 1 ml column (GE Healthcare) charged with Ni^2+^ and attached to an AKTA system. The column was washed with 25 mM sodium phosphate buffer pH 7.5, 500 mM NaCl, 10 % glycerol, 0.1% NP-40, 25 mM imidazole, 1 mM DTT, 0.25 mM PMSF. Recombinant Sgo1 protein was eluted with an increasing imidazole gradient (25 - 500 mM) over 40 column volumes and then loaded onto a gel filtration Superose 6 10/300 column in 50 mM Tris-HCl pH 7.5, 500 mM NaCl, 10% glycerol, 0.1 mM EDTA, 0.1 mM EGTA, 1 mM DTT, 0.25 mM PMSF. Fractions containing the eluted protein were run on an SDS-PAGE gel and the purity of recombinant Sgo1 was assessed by InstantBlue (Expedeon) staining.

### Cloning, expression and purification of of SUMOylation enzymes

SUMO E1 enzymes Uba2 and Aos1 (from vectors pET11a-*UBA2* and pET28a-*AOS1*, [53]) were co-expressed in BL21 CodonPlus (DE3) RIL (Agilent Technologies) in the presence of 0.05 mM IPTG at 18 °C for 20 h. Expression of SUMO E2 enzyme Ubc9 and SUMO Smt3 (pET21b-*UBC9* and pET21a-*SMT3* [53]) was induced by 0.1 mM IPTG at 18 °C for 20 h in BL21 CodonPlus (DE3) RIL. E1, E2 and SUMO were purified using Ni-NTA agarose resin (Qiagen) with 20 mM sodium phosphate pH7.5, 500 mM NaCl, 10 to 200 mM imidazole, followed by Superdex 200 16/600 column (hand poured using prep grade resin from GE Healthcare) in 50 mM Tris-HCl pH 7.5, 150mM NaCl, 1mM DTT, 10% glycerol.

*siz1-167-508* and *siz2-130-490* were amplified from plasmids pGEX-4T-1-*SIZ1-HF* [53]) and AMp1528 (YEPlac112-*SIZ2*, this work), respectively. The respective PCR product was cloned into pMAL-c2X to generate N-terminal MBP-tagged *siz1-167-508* (AMp1679) and N-terminal MBP-tagged *siz2-130-490* (AMp1680). Siz1 (167-508) or Siz2 (130-490) were expressed in LB containing 10 μM ZnCl_2_, 0.1 mM IPTG and appropriate antibiotics at 16 °C for 20 h using BL21 CodonPlus (DE3) RIL (Agilent Technologies). Siz1 (167-508) or Siz2 (130-490) was purified from MBPTrap HP 1 mL column (GE Healthcare) with 50 mM Tris-HCl pH 7.5, 150 mM NaCl, 10 μM ZnCl_2_, 0 to 10 mM maltose gradient, followed by gel filtration with a Superdex 200 16/600 column (hand poured using prep grade resin from GE Healthcare) in 50 mM Tris-HCl pH 7.5, 150 mM NaCl, 10 μM ZnCl_2_, 10% glycerol.

### *In vitro* SUMOylation assays and SUMOylated Sgo1 pull down assays

Purified Sgo1 (1 μM) was mixed with 5 mM ATP, 15 μM SUMO, 0.5 μM E1 (unless otherwise indicated), 0.5 μM E2 (unless otherwise indicated), 0.1 μM (unless otherwise indicated) Siz1 (167-508) or Siz2 in a reaction buffer consisting of 25 mM HEPES pH 7.5, 150 mM KCl, 10 mM MgCl_2_, 15 % glycerol, 0.1 % NP-40, 0.1 mM DTT, 0.25 mM PMSF. The reaction was incubated at 30 °C for 2 h (unless otherwise indicated). For detection of SUMOylated Sgo1, the reaction was boiled in SDS sample buffer before analyzed by anti-V5 western blotting.

For the binding assay, product of *in vitro* SUMOylation was incubated with 50 μL of anti-V5 coupled protein G magnetic dynabeads (Thermofisher) in Binding buffer A (25 mM HEPES pH 7.5, 150 mM KCl, 2 mM MgCl_2_, 15 % glycerol, 0.1% NP-40, 0.1 mM EDTA, 0.5 mM EGTA, 0.25 mM PMSF) for 2.5 h at 4 °C. After washing three times with Binding buffer A, beads were incubated with yeast lysate containing tagged Rts1 or Bir1 for 2 h at 4 °C. Beads were washed five times with binding buffer A + 10 mM NEM and were heated at 65 °C for 15 min to elute bound proteins.

### Analysis of *in vivo* ubiquitination

Cultures were inoculated to OD_600_ = 0.2 in 100 mL 2% raffinose-containing dropout media and were arrested in G1 as described above. 2% galactose was added for 30 min to induce *pGAL-SGO1-9MYC* expression. Cells were treated, lysed in buffer A and diluted as described in *in vivo* SUMOylation assay. Lysates were incubated with Ni-NTA beads, 0.05% Tween-20 and 15 mM imidazole overnight at 4°C, with gentle rotation. Beads were washed twice with buffer A + 0.05% Tween-20, and four times with buffer C (8M urea, 100 mM sodium phosphate buffer pH 6.3, 10 mM Tris-HCl, pH 6.3) + 0.05% Tween-20. For elution, beads were heated at 60°C for 10 min in HU buffer (8M urea, 200 mM Tris-HCl pH 6.8, 1 mM EDTA, 5% (w/v) SDS, 0.1% (w/v) bromophenol blue, 1.5% (w/v) DTT).

### Sgo1 half-life measurement

Cultures were inoculated to OD_600_ = 0.2 in YEP + 2% raffinose + 0.3 mM adenine. 5 μg/mL alpha-factor was added for 1.5 h and re-added to 2.5 μg/mL every hour until the end of the experiment. *SGO1* overexpression was induced by the addition of 2% galactose for 30 min. *De novo* synthesis of Sgo1 protein was quenched by the addition of 2% glucose and 1 mg/mL cycloheximide (Acros Organics).

### Co-immunoprecipitation

Cultures were inoculated to OD_600_ = 0.2 in 2 L YPDA and grown at 30 °C until OD_600_ reached 1-1.5. Cell pellets were harvested by centrifugation and drop-frozen in liquid nitrogen. Cells were ground, lysed and benzonase-treated as described in ‘Purification of recombinant protein’. After centrifugation at 3,600 rpm for 10 min, protein concentrations were measured by Bradford assay and approximately 85 mg of protein was incubated with 4 mg of epoxy beads (Thermofisher) coupled to IgG for 1.5 h at 4°C. Beads were washed four times with the lysis buffer and were heated at 70°C for 10 min in 50 μL 1 × LDS sample buffer + 2.5% β-mercaptoethanol. 5 μL of input and 20 μL of IP samples were separated on an 8% acrylamide gel and were transferred to PVDF membranes. Western blot was performed using TBST-based buffer.

### Chromatin immunoprecipitation (ChIP)

Metaphase-arrested cells were fixed in 1.1% formaldehyde for at least 30 min. Cells were washed twice with TBS (20 mM Tris-HCl pH 7.5, 150 mM NaCl), once with FA lysis buffer (50 mM HEPES-KOH pH 7.5, 150 mM NaCl, 1 mM EDTA, 1% Triton-X, 0.1% sodium deoxycholate) + 0.1% SDS, and drop frozen in liquid nitrogen. Cells were lysed by bead beating in FA lysis buffer + 0.5% SDS + 1 mM PMSF + EDTA-free protease inhibitor cocktail (Roche). The resulting pellets were sonicated in a Bioruptor machine. Sheared chromatin was incubated with the relevant antibody and protein G dynabeads (Thermofisher) at 4°C overnight. Anti-HA (12CA5, Roche Diagnostics), anti-Flag M2 (Sigma), or anti-V5 (Abd Serotec) antibodies were used as relevant. Beads were washed once with FA lysis buffer + 0.1% SDS+ 275 mM NaCl, once with FA lysis buffer + 0.1% SDS + 500 mM NaCl, once with wash buffer (10 mM Tris-HCl pH 8, 0.25 M LiCl, 1 mM EDTA, 0.5% NP-40, 0.5% sodium deoxycholate) and once with TE (10 mM Tris-HCl, pH 8, 1 mM EDTA). Chromatin was eluted by boiling in the presence of 10% Chelex beads, treated with proteinase K (Life technologies) for 30 min and boiling for a further 10 min. Quantitative PCR was done using Express SybrGreenER (Thermofisher) (*CEN4* and *ARM4*) or Luna (NEB) (all other chromosomal sites) on a Lightcycler machine (Roche). Primers used were listed in Table S5. ChIP enrichment was determined using the formula E^-ΔCt^, where ΔCt = (Ct_(ChIP)_ – [Ct_(input)_ – logE(Input dilution factor)] and E represents primer efficiency.

### Biorientation assay in fixed and live cells

Cells carrying *pMET-CDC20*, *SPC42-tdTOMATO* and *CEN4-GFP* were arrested in metaphase in the presence of nocodazole and benomyl as described in ‘yeast growth and synchronization’.

For analyzing biorientation in metaphase-arrested cells (Figure 5C and 5D), drugs were washed out by filtering with 10 times the volume of YEP and cells were released into YPDA + 8 mM methionine. Samples were taken at the indicated time points, fixed in 3.7% formaldehyde for 9 min, washed in 80% ethanol and resuspended in 1 μg/mL DAPI in PBS. 200 cells were analyzed for each time point using fluorescent microscope.

For analyzing biorientation in cells going into anaphase (Figure 5E and 7F), nocodazole-arrested cells were loaded onto the ONIX Microfluidic Perfusion System (CellAsic) and visualized with a Zeiss inverted microscope coupled to an EMCCD camera at 25°C. Imaging started as soon as cells were released into methionine dropout media without drugs.

## Supplementary information

**Table S1.** Complete list of high copy suppressors of *GAL-SGO1* sickness identified in the screen shown in Figure S1A.

**Table S2.** (A) List of SUMOylated peptides identified in mass spectrometry (B) List of peptides identified in mass spectrometry

**Table S3.** Yeast strains used in this study

**Table S4.** Plasmids used in this study

**Table S5.** Oligonucleotides used in this study

---

**Figure S1.**
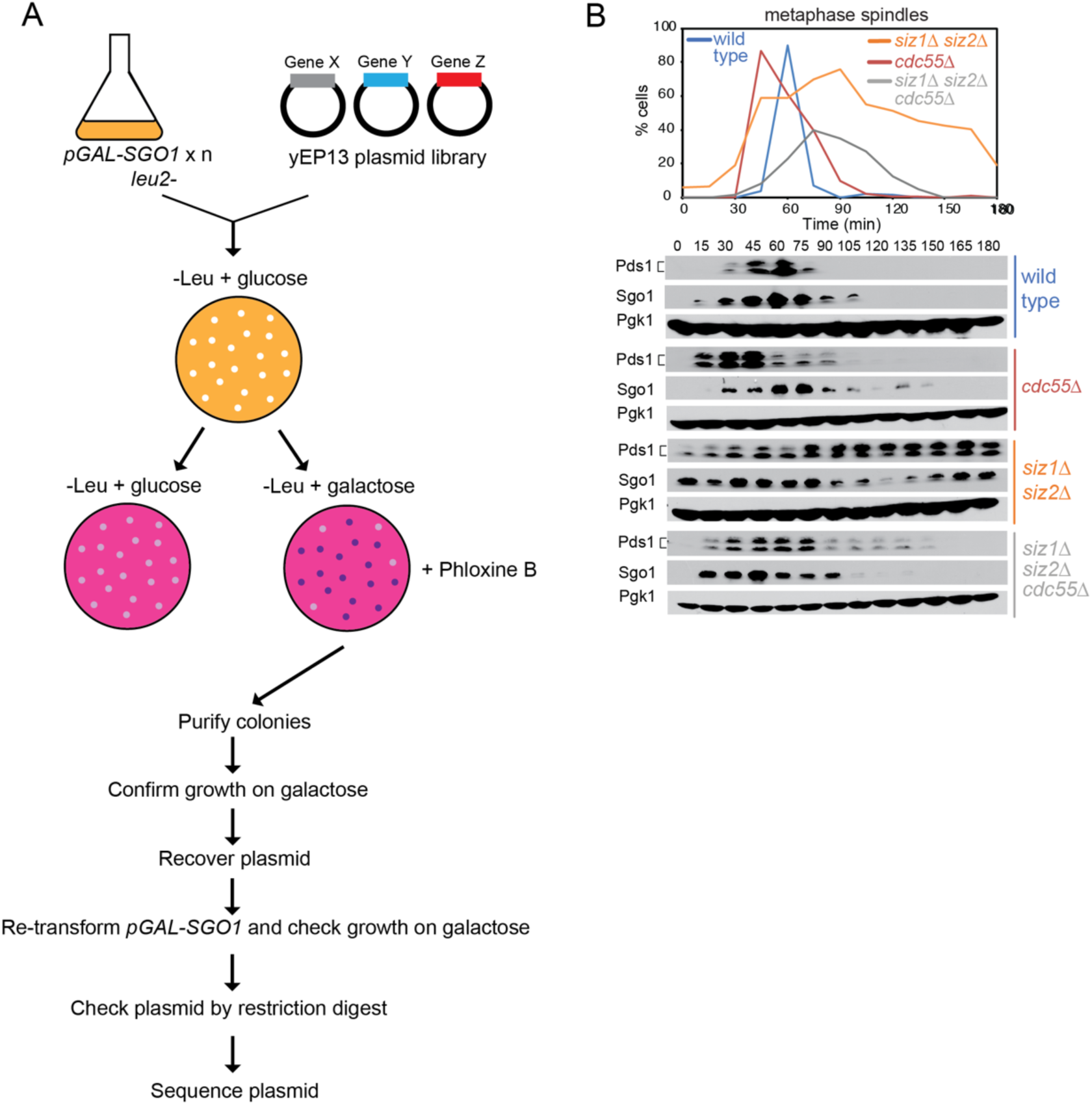
**related to Figure 1**. Identification of SUMO ligases as Sgo1 regulators. (A) Overview of multi-copy suppressor screen to identify Sgo1 antagonists. (B) Deletion of *CDC55* partially alleviates the metaphase delay phenotype of the *siz1*Δ *siz2*Δ mutant. Mitotic time course analysis was performed as described in Figure 1F, for the following strains: wild type (AMy8467), *cdc55Δ* (AMy8779), *siz1*Δ *siz2*Δ (AMy8452) and *siz1*Δ *siz2*Δ *cdc55Δ* (AMy8637).

**Figure S2.**
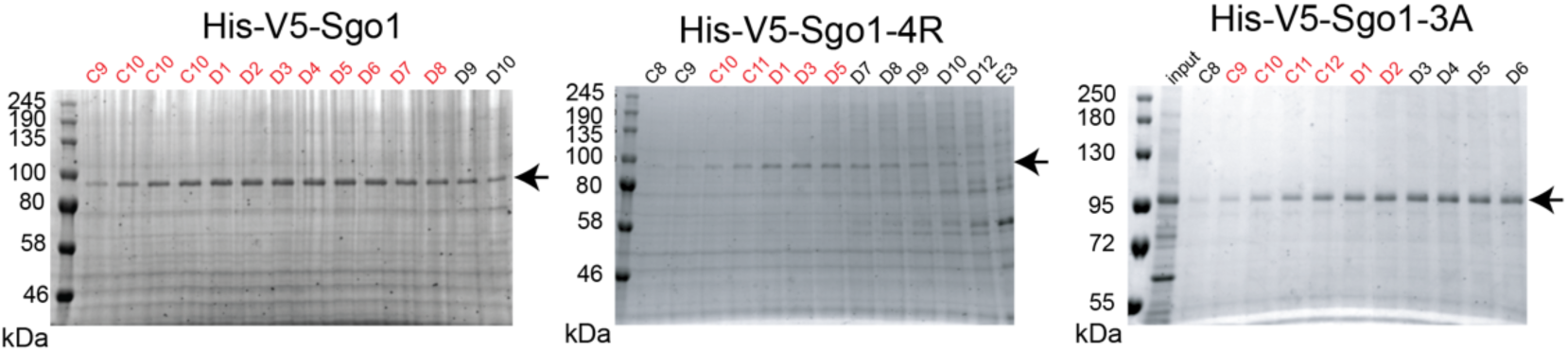
**related to Figure 2**. Purification of Sgo1. Coomassie staining confirmed the successful purification of wild type and mutant Sgo1 recombinant proteins.

**Figure S3.**
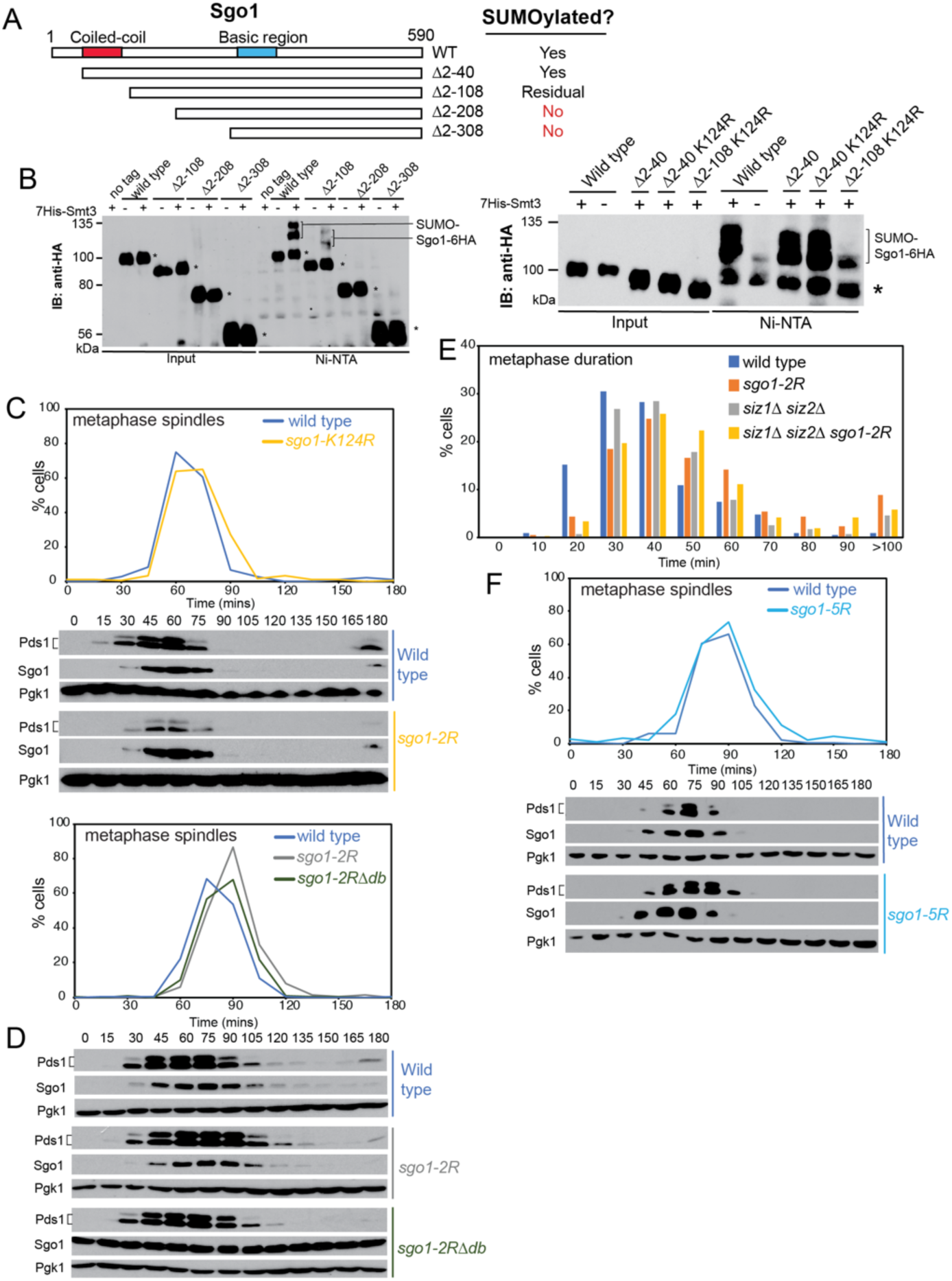
**related to Figure 3**. Identification and characterization of unSUMOylatable Sgo1 mutants. (A) Schematics describing the truncation mutants generated for Sgo1. The conserved coiled-coil and basic domains are highlighted in red and blue, respectively. Results from (B) are summarized on the right. (B) Sgo1 is likely to be SUMOylated in the first 208 amino acids. *In vivo* SUMOylation was assessed for the following Sgo1-6HA tagged strains as described in Figure 2A, together with the indicated negative controls: wild type (AMy7654), *sgo1Δ2-108* (AMy14764), *sgo1*Δ *2-208* (AMy14765) and *sgo1Δ2-308* (AMy14766). Unmodified Sgo1 bands are marked with asterisks. In addition to Lys124 SUMOylation, identified by mass spectrometry, the region between amino acids 41 and 108 is likely to be SUMOylated. *In vivo* SUMOylation was assessed for the following Sgo1-6HA tagged strains: wild type (AMy7654), *sgo1*Δ*2-40* (AMy18194), *sgo1*Δ*2-40 K124R* (AMy18476) and *sgo1*Δ*2-108 K124R* (AMy16540). (C-F) Characterization of unSUMOylatable Sgo1 mutants. (C) The Sgo1-K124R mutants does not show a metaphase delay. Mitotic time course analysis as described in Figure 1F was performed for wild type (AMy8467) and *sgo1-K124R* (AMy24448) strains carrying *SGO1-9MYC* and *PDS1-HA*. (D) The *sgo1-2R* mutant shows a very minor metaphase delay that is not exacerbated by mutations in the Sgo1 destruction box. Analysis of *sgo1-2R* (AMy24448) and *sgo1-2R-*Δ*db* (AMy25924) cells carrying *SGO1-9MYC* and *PDS1-3HA* as described in Figure 1F. (E) Metaphase duration was measured after live cell imaging of wild type (AMy24546), *siz1*Δ *siz2*Δ (AMy24547), *sgo1-2R* (AMy24573) and *sgo1-2R siz1*Δ *siz2*Δ (AMy24575) strains carrying *YFP-TUB1* and *CDC14-GFP* as described in Figure 1D. (F) *sgo1-5R* (AMy24450) does not show a metaphase delay, suggesting adverse effects of K124R on Sgo1 protein function when combined with the 4R mutations.

**Figure S4.**
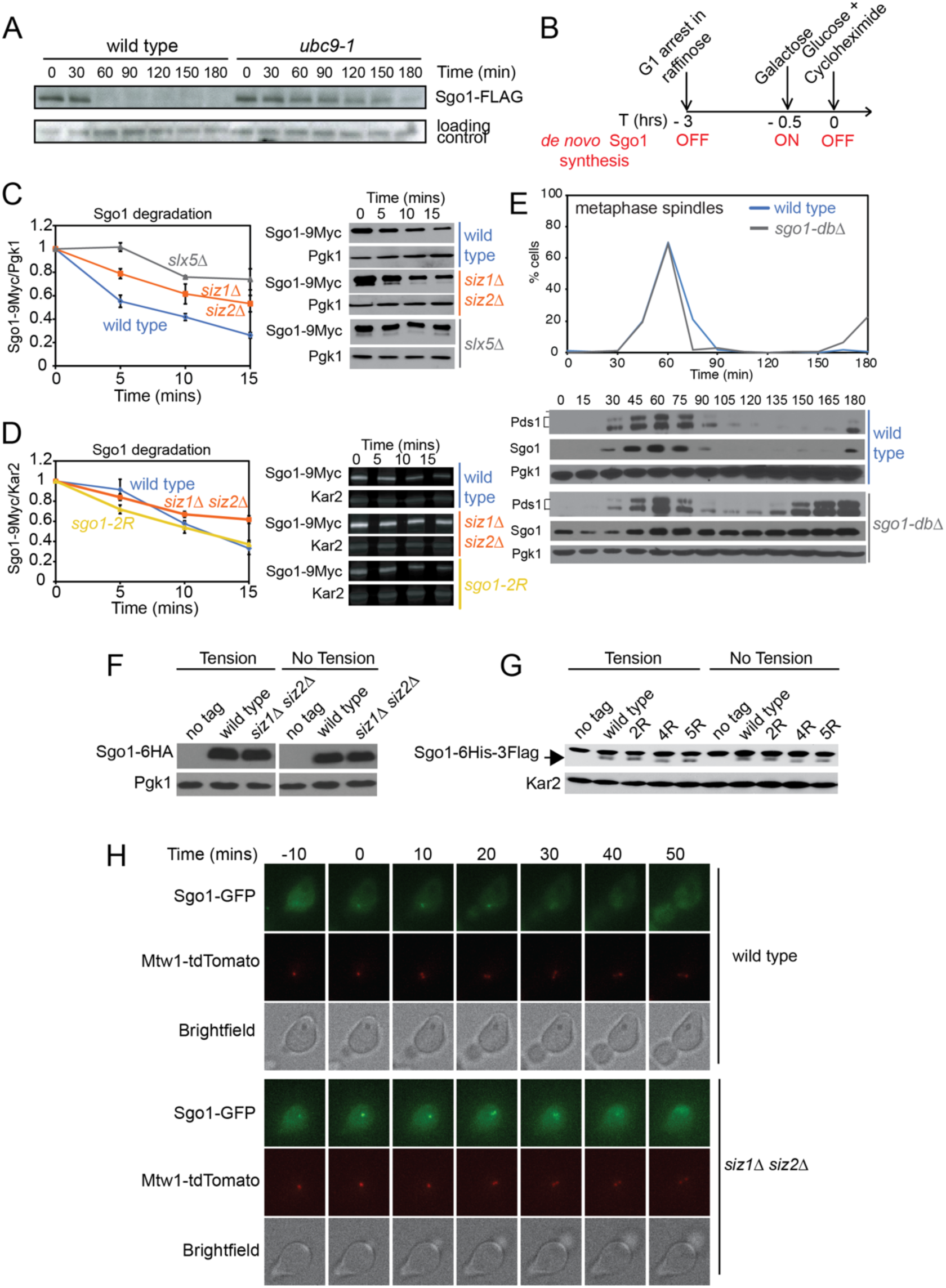
**related to Figure 4**. Analysis of factors promoting Sgo1 degradation. (A) The degradation of Sgo1 depends on SUMO-conjugating protein Ubc9. The cells were synchronized in nocodazole and released into medium with α-factor to ensure arrest in G1. (B and C) Sgo1 half-life is increased in *slx5*Δ and *siz1*Δ *siz2*Δ mutants. (B) Scheme describing the cycloheximide chase experiment. Cells were arrested in G1 throughout the experiment and *pGAL1-SGO1-9MYC* expression was initially prevented by growth of cells in raffinose. Subsequently, a pulse of Sgo1 was provided by the addition of galactose, after which *de novo* Sgo1-9Myc synthesis was quenched by the addition of glucose (to block *pGAL* expression) and cycloheximide (to block protein synthesis). (C) Cycloheximide chase experiment was performed for *pGAL-SGO1-9MYC* (AMy1392), *siz1*Δ *siz2*Δ *pGAL-SGO1-9MYC* (AMy22584), and *slx5*Δ *pGAL-SGO1-9MYC* (AMy22586) strains. Representative anti-Myc immunoblot (right) to detect Sgo1-9Myc is shown. Pgk1 is shown as a loading control. Quantification showing the mean values from three biological replicates (left) is also shown. Relative Sgo1 levels were calculated as the ratio of Myc signal to Pgk1 signal and the ratio was set to 1 for time 0. Error bars represent standard error calculated from three independent experiments. (D) The effects of SUMOylation on Sgo1 half-life are likely to be indirect. Sgo1 degradation is unaffected in the *sgo1-2R* mutants. Cycloheximide chase experiment was performed for *pGAL-SGO1-9MYC* (AMy1392), *siz1*Δ *siz2*Δ *pGAL-SGO1-9MYC* (AMy22584) and *pGAL-sgo1-2R-9MYC* (AMy24940) strains. Kar2 was probed as a loading control. Quantification and statistics were performed as described in (C). (E) Sgo1 degradation is not required for timely anaphase entry. Time course analysis of wild type (AMy18500) and *sgo1-*Δdb (AMy18440) strains carrying *SGO1-6HA* and *PDS1-18MYC* as described in Figure 1F. (F-H) SUMOylation does not regulate Sgo1 dissociation from pericentromeres in response to tension. (F) Sgo1 expression is unchanged in metaphase-arrested *siz1*Δ *siz2*Δ cells. Protein extracts from Figure 4F were analysed by anti-HA and anti-Pgk1 (loading control) western blotting. (G) Sgo1 expression is unchanged in metaphase-arrested *sgo1-2R, 4R and 5R* mutants. Protein extracts from Figure 4G were analysed by anti-Flag and anti-Kar2 (loading control) western blotting. The non-specific bands were marked with an asterisk. (H) Sgo1 localization in metaphase was unchanged in the *siz1*Δ *siz2*Δ cells. Wild type (AMy9233) and *siz1*Δ *siz2*Δ (AMy15604) strains carry *pMET-CDC20*, *SGO1-yeGFP* and *MTW1-tdTomato*. Shown are representative pictures of cells. The point at which a bright Sgo1 focus was formed was assigned as time 0.

**Figure S5.**
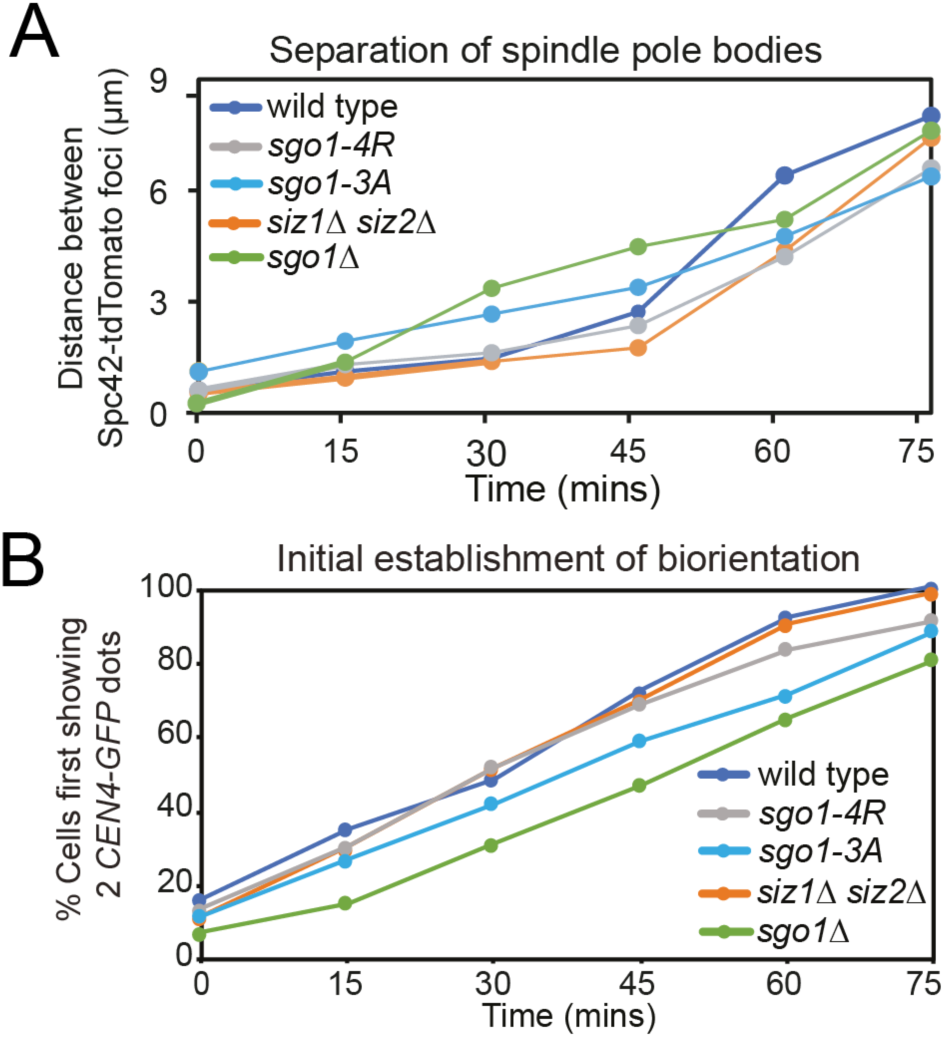
**related to Figure 5**. Sister kinetochore biorientation is proficient in SUMO mutants, but not in *sgo1*□ or *sgo1-3A*. Spindle pole body separation (A) and initial establishment of biorientation for the experiment shown in Figure 5C-D. (A) The distance between Spc42-tdTomato dots was measured by ImageJ and the average distance was calculated for each time point. (B) The initial establishment of biorientation is unaffected in *siz1*Δ *siz2*Δ and *sgo1-4R* cells. The time point at which a cell first displayed two *CEN*4-GFP dots was defined as the timing of the initial establishment of biorientation.

**Figure S6.**
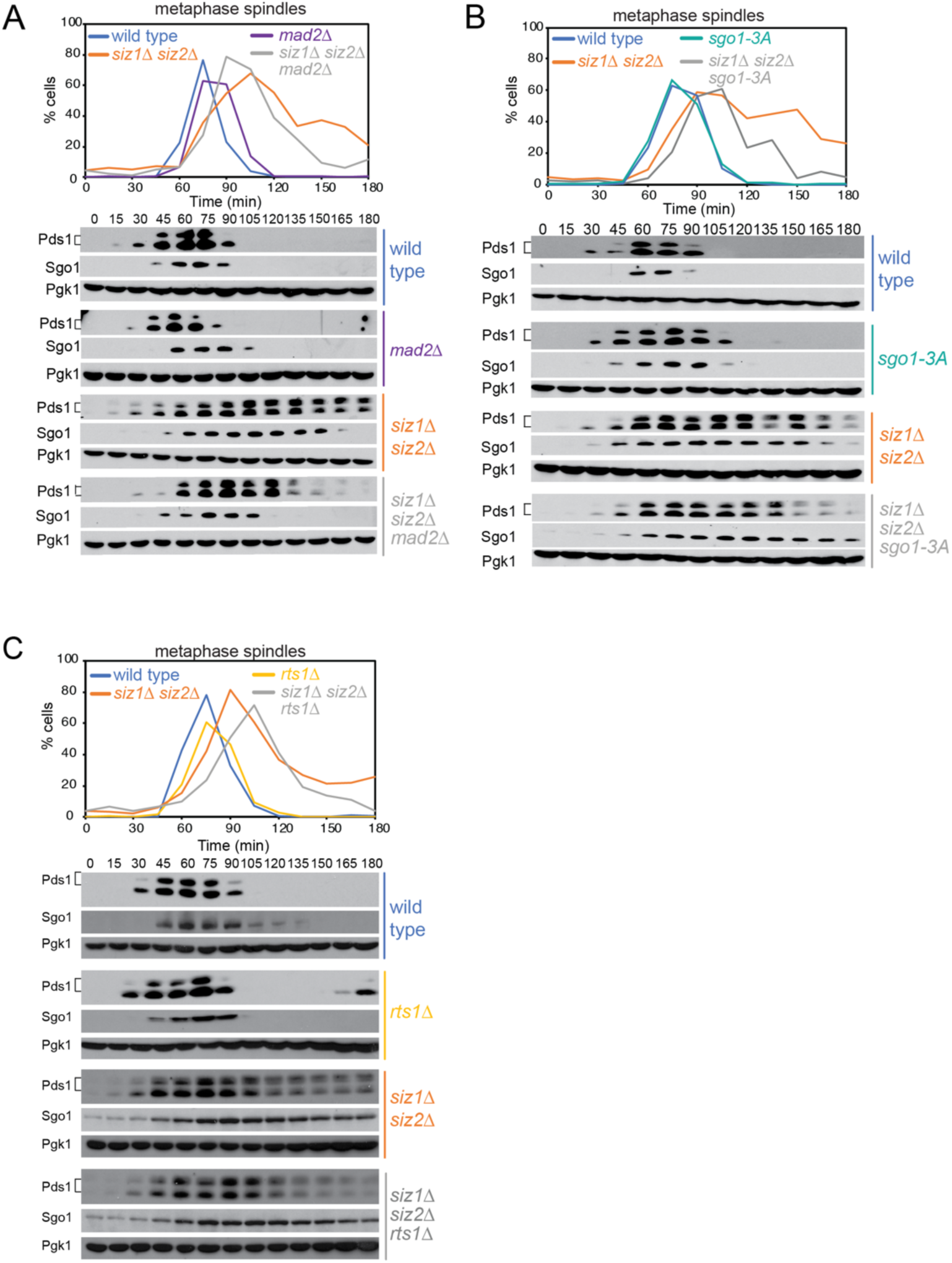
**related to Figure 6.** Inactivation of CPC and SAC, but not PP2A-Rts1, rescues the metaphase delay of *siz1*Δ *siz2*Δ cells. (A) Deletion of the SAC component, *MAD2*, partially rescued the metaphase delay of *siz1*Δ *siz2*Δ mutant. Mitotic time course was performed as in (A) for the following strains: wild type (AMy8467), *mad2Δ* (AMy9635), *siz1*Δ *siz2*Δ (AMy8452) and *siz1*Δ *siz2*Δ *mad2*Δ (AMy9634). (B) Partial rescue of the metaphase delay of *siz1*Δ *siz2*Δ cells by the *sgo1-3A* mutation. Wild type (AMy8467), *siz1*Δ *siz2*Δ (AMy8452), *sgo1-3A* (AMy8964) and *siz1*Δ *siz2*Δ *sgo1-3A* (AMy8755) strains carrying *PDS1-HA* and *SGO1-9MYC* were analysed as described in Figure 1F. (C) Deletion of *RTS1* did not rescue the metaphase delay of the *siz1*Δ *siz2*Δ mutant. Mitotic time course was performed as in (A) for the following strains: wild type (AMy8467), *rts1*Δ (AMy20909), *siz1*Δ *siz2*Δ (AMy8452) and *siz1*Δ *siz2*Δ *rts1*Δ (AMy17284).

**Figure S7.**
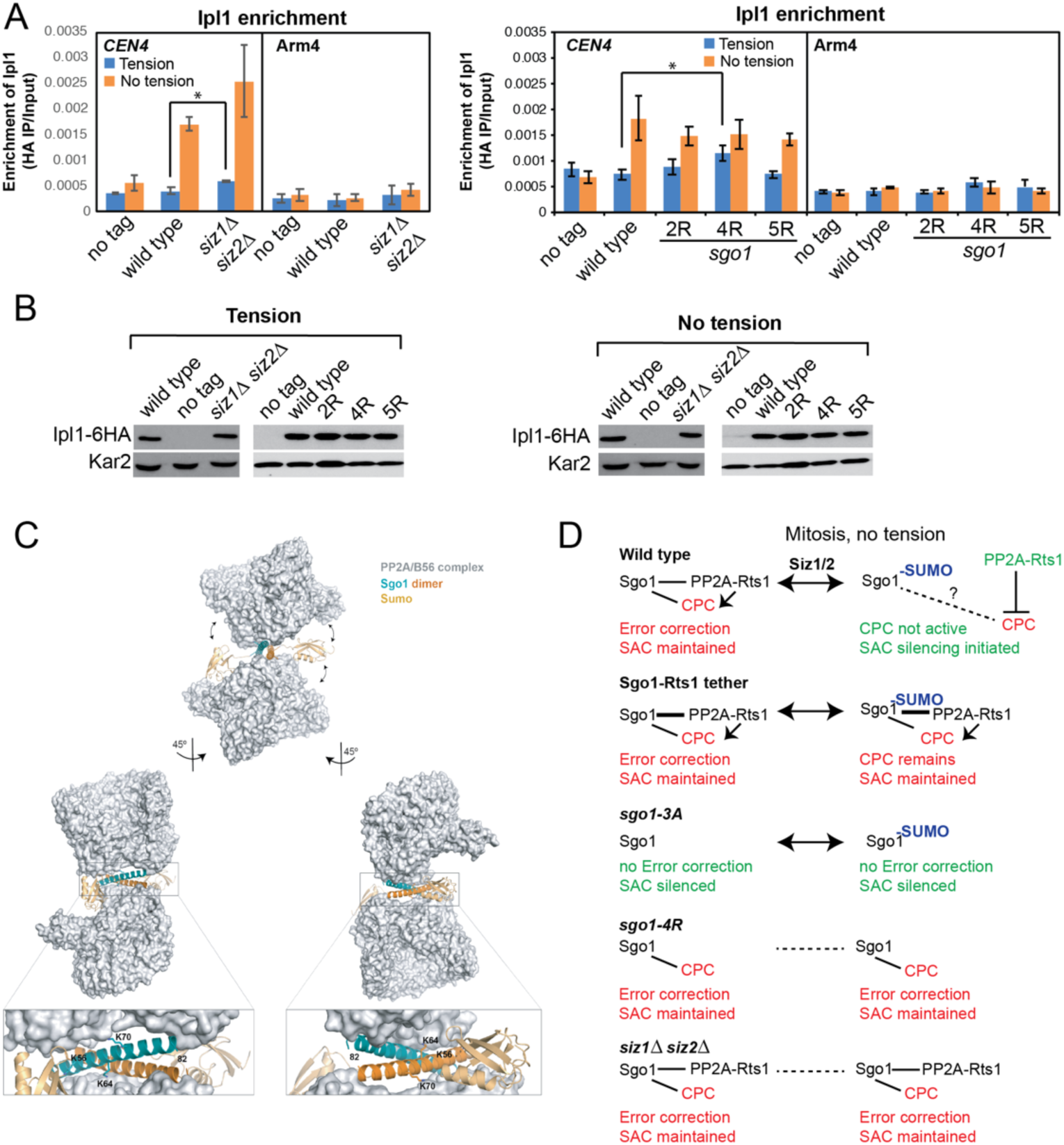
**related to Figure 7.** In Sgo1 SUMO-deficient cells, Ipl1 is not completely removed when the cells are under tension. (A) Ipl1 association with *CEN4* and *ARM4* were measured by ChIP-qPCR using wild type (AMy26686), *siz1*Δ *siz2*Δ (AMy23194), *sgo1-2R* (AMy26684), *sgo1-4R* (AMy26692) and *sgo1-5R* (AMy26691) carrying *IPL1-6HA*, together with a no tag control (AMy2508). Cells were arrested in metaphase by depletion of Cdc20 in the presence or absence of spindle tension. Error bars represent standard errors calculated from 5 biological repeats. * = *P* < 0.05. (B) Ipl1 protein levels are unchanged in Sgo1 SUMO-deficient mutants. Protein extracts from (A) were analyzed by anti-HA and anti-Kar2 (loading control) western blotting. (C) Structural modelling predicts that SUMOylation on the coiled-coil domain of Sgo1 is incompatible with Sgo1-PP2A interaction. *S.c*. Sgo1-PP2A interaction was modelled based on structural information obtained from co-crystallized human Sgo1(51-96) and PP2A using Phyre2 web portal (www.sbg.bio.ic.ac.uk/phyre2) [54]. Potential consequence of symoylation was modelled using the molecular graphic program PyMOL (The PyMOL Molecular Graphics System, Version 2.0 Schrödinger, LLC). According to this model, Lys64 and Lys70 are critically positioned at the binding surface with no room to accommodate a bulkier modification such as sumoylation. Lys56 is exposed to the solvent, but the attachment of SUMO (highlighted in gold) is expected to result in steric clashes with PP2A and weaken Sgo1-PP2A binding. Structural information is unavailable beyond Leu82 and so Lys85 could not be included in this model. (D) Model for role of Sgo1 SUMOylation in stabilizing the bioriented state. In **wild type cells**, Sgo1 brings both PP2A-Rts1 and CPC to centromeres and PP2A-Rts1 enhances CPC localization. A minor pool of Sgo1 is dynamically SUMOylated and this both prevents PP2A-Rts1 binding and directly or indirectly promotes CPC removal, dampening its activity at kinetochores. Upon **tethering of PP2A-Rts1 to Sgo1**, release of PP2A-Rts1 cannot occur and CPC activity persists. In *sgo1-3A* cells, the interaction with both PP2A-Rts1 and CPC is absent and error correction is defective due to a failure to maintain CPC. In ***sgo1-4R*** cells, PP2A-Rts1, but not CPC binding is lost. The failure to SUMOylate, along with the absence of PP2A-Rts1, means that the phosphatase cannot undergo its capture and release by Sgo1. Either as a consequence of this and/or other effects of Sgo1 SUMOylation, CPC is maintained on kinetochores resulting in ectopic error correction and destabilization of kinetochore-microtubule attachments. ***siz1*Δ *siz2*Δ** mutants show similar behaviour to *sgo1-4R* except that Sgo1 is expected to retain the ability to bind PP2A-Rts1, so that absence of SUMOylation both prevents CPC removal and PP2A-Rts1 release.

